# Scinderin-driven Golgi Actin Remodeling coordinates GLP-1 and insulin secretion to regulate glucose homeostasis

**DOI:** 10.64898/2026.05.28.728433

**Authors:** Laura Martin-Diaz, Maarit S. Patrikainen, Chen Chongtham, Alka Gupta, Valeriia Dotsenko, Markus J.T. Ojanen, Fábio Tadeu Arrojo Martins, Joel Johnson George, Roselia Davidsson, Satu Hakanen, Kim Eerola, Eriika Savontaus, Jonna Saarimäki-Vire, Diego Balboa, Jorma Isola, Jutta E. Laiho, Heikki Hyöty, Timo Otonkoski, Maija Vihinen-Ranta, Diana M. Toivola, Keijo Viiri

**Author notes:** Correspondence. Address correspondence to: Keijo Viiri, PhD, Faculty of Medicine and Health Technology, Tampere University Hospital, Tampere University, Arvo Ylpön Katu 34, Tampere, FIN-33520, Finland.

## Abstract

Maintenance of glucose homeostasis requires coordinated hormone secretion from intestinal enteroendocrine cells and pancreatic β-cells, yet the intracellular mechanisms that couple nutrient sensing to endocrine output remain poorly defined. Here, we identify the actin remodeler Scinderin (SCIN) as a shared regulator of hormone secretion across these systems. SCIN is selectively expressed in enteroendocrine L-cells and pancreatic β-cells, where it localizes to phosphatidylinositol-4-phosphate (PI(4)P)-enriched Golgi membranes and controls Golgi-associated actin dynamics. Loss of SCIN disrupts Golgi organization, impairs prohormone trafficking, and reduces secretory granule formation, resulting in defective nutrient-stimulated GLP-1 and insulin secretion while preserving cAMP-dependent amplification pathways. In vivo, tissue-specific deletion of Scin compromises incretin responses, β-cell insulin secretion, and systemic glucose homeostasis. Consistent with these findings, SCIN expression is reduced in human diabetic β-cells and associates with stress-related loss of β-cell maturity. Transcriptomic analyses reveal a conserved Golgi stress program upon SCIN loss, linking intracellular trafficking defects to endocrine dysfunction. Together, our results identify SCIN-dependent Golgi actin remodeling as a rate-limiting intracellular mechanism coordinating enteroendocrine and pancreatic hormone secretion. This work uncovers a shared, targetable node controlling endocrine output, providing a mechanistic link between secretory pathway dysfunction and diabetes.

## Main

Endocrine cells in the intestine and pancreas maintain metabolic homeostasis by translating nutrient cues into tightly regulated hormone secretion^1,2^. This task places exceptional demands on the Golgi apparatus, where prohormones are sorted, modified and packaged into dense-core granules^3,4^. Increasing evidence highlights the Golgi-associated F-actin network as an active mechanical scaffold that shapes membrane curvature, drives carrier budding and supports granule maturation^5^. Contrary to our detailed understanding of cortical actin regulation^6–8^, the factors specifying lineage-specific, nutrient-responsive Golgi F-actin dynamics remain elusive.

Our previous and current work and developmental epigenomic datasets identify Scinderin (SCIN) as an unusually selective component of this landscape. Unlike many ubiquitous actin regulators, *SCIN* is developmentally regulated by Polycomb Repressive complex 2 (PRC2) in both intestine^9^ and pancreas^10^: it is silenced in intestinal stem-state epithelium and derepressed upon differentiation, and emerges late in β-cell ontogeny in concert with the opening of L-cell– and β-cell–restricted enhancers at the *SCIN* locus. This pattern situates SCIN at the interface of developmental identity and secretory competence, suggesting that SCIN may endow mature endocrine cells with a specialized Golgi-actin architecture calibrated for high-output hormone production.

This mechanistic question carries particular relevance for diabetes, a disease in which endocrine failure extends beyond insulin deficiency to encompass defects in prohormone processing, Golgi stress, granule maturation, and disrupted enteroendocrine–pancreatic coupling^2,11^. While current therapies improve glycaemic control, they largely act downstream of the intracellular pathways governing hormone biosynthesis, granule maturation, and nutrient-stimulated secretion^12^. Understanding how lineage-restricted actin regulators such as SCIN integrate developmental programs with Golgi function may therefore uncover root-cause vulnerabilities shared across L-cells and β-cells and open therapeutic avenues not addressed by existing treatments.

## Results

### Scinderin is selectively expressed and developmentally controlled in endocrine cells that regulate glucose homeostasis

Our previous work identified *Scin* as a PRC2 target gene in the mouse intestine^9^ and we began to analyse the expression of *Scin* in intestine and pancreas tissues. SCIN protein was highest in differentiated (ENRI) intestinal organoids, placing *Scin* in villus lineages rather than stem cells (WENRC), and qPCR in mouse intestinal organoids showed *Scin* enrichment under goblet-(ENRID) and enteroendocrine-enriched (ENRID+iMEK) conditions (Fig. 1a; Extended Data Fig. 1a–b). In mouse intestine, SCIN colocalized with GLP-1^+^ L-cells, with additional co-occurrence with PYY, but not with 5-HT, SST, GIP or CCK (Fig. 1b; Extended Data Fig. 1f). Single-cell datasets further localized *Scin/SCIN* to distal small intestine and to enteroendocrine L-cells: expression was highest in ileum and enriched within glucagon-expressing clusters in both mouse and human organoid-derived enteroendocrine cells (EECs) (Fig. 1c; Extended Data Fig. 1c–e).

**Figure 1.**
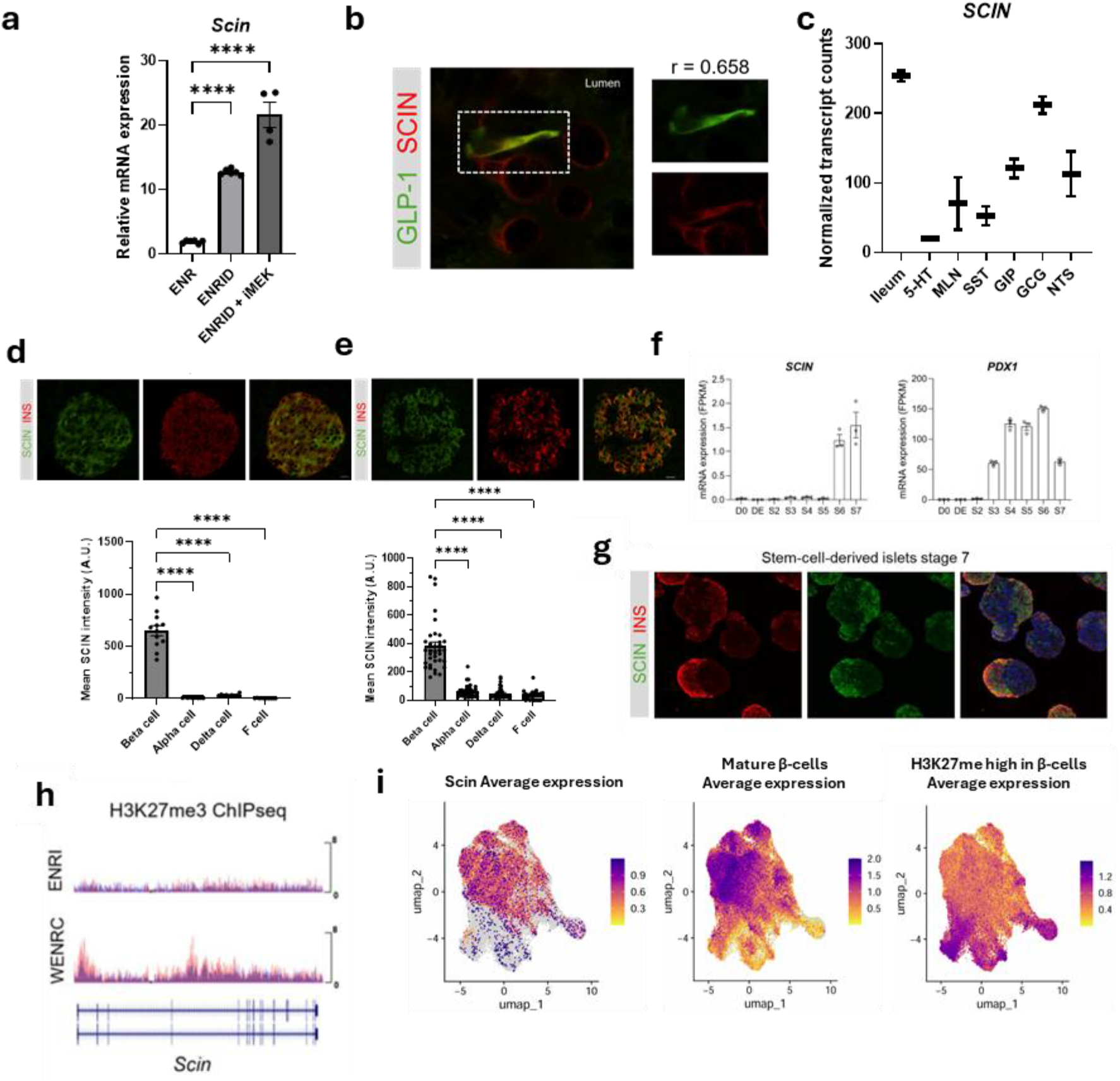
Scinderin is selectively expressed and developmentally regulated in intestinal L-cells and pancreatic β-cells. **a**, qPCR shows *Scin* enrichment in goblet- (ENRID) and EEC (ENRID+iMEK)-enriched mouse organoids versus control (ENR). Data are presented as mean ± SEM. ****p ≤ 0.0001 vs. indicated groups by one-way ANOVA with Dunnett’s post-hoc test (n = 4– 6). **b**, SCIN colocalizes with GLP-1^+^ L-cells in mouse intestine. R = Pearson’s correlation coefficient (n = 4) **c**, Human ileal scRNAseq and -organoid derived-EECs^14^ (n = 2 independent experiments) show highest SCIN expression in GCG^+^ (GLP-1) populations. Data are presented as median with min-max values. **d, e**, (Top) In mouse and human pancreas, SCIN is predominantly detected in insulin (INS) producing-β-cells. Scale bar, 20 µm. (Bottom) Quantification of SCIN intensity in hormone-identified islet cells (n = 8–12 islets from 3 mice and n = 24–35 islets from 6 human donors). Data are presented as mean ± SEM. ****p ≤ 0.0001 vs. indicated groups by one-way ANOVA with Dunnett’s post-hoc test. **f**, SCIN and PDX1 expression throughout *in vitro* stem cell differentiation into functional SC-islets determined by bulk RNA sequencing^15^. SCIN emerges at late stages of differentiation, PDX1 used as marker for β-cell development. Representative stages of the pancreatic-directed differentiation are presented: stem cells (D0), definitive endoderm (DE), primitive gut tube (S2), posterior foregut (S3), pancreatic progenitors (S4), endocrine progenitors (S5), immature SC-islets (S6) and functional mature SC-islets (S7). Values are presented as fragments per kilobase million (FPKM) (n = 3 independent experiments). **g**, SCIN colocalization with INS expressing β-cells from stem-cell-derived islets. **h**, H3K27me3 ChIP–seq indicates PRC2-dependent repression of Scin in -stemstate- (WENRC) intestinal organoids, relieved upon differentiation (ENRI)^9^. **i**, scRNAseq analysis^16^ reveals *SCIN* is expressed in the mature β-cells subpopulation but is reduced in the high H3K27me3 β-cell subpopulation.

In the pancreas, SCIN was predominantly detected in β-cells in both mouse and human islets by immunostaining and was supported by islet RNA-seq and chromatin accessibility at the *SCIN* locus (Fig. 1d, e; Extended Data Fig. 2a–d). During human stem-cell–derived islet differentiation, *SCIN* expression emerged at late stages and colocalized with insulin^+^ cells (Fig. 1f-g), paralleling developmental data from human fetal pancreas and epigenomic profiling that revealed progressive promoter accessibility and intestinal and β-cell–specific upstream enhancers bound by PDX1, NKX2-2 and NKX6-1 (Extended Data Fig. 3a–d).

Epigenetically, PRC2 constrained *Scin* in intestinal stem-state organoids (H3K27me3 in WENRC) with derepression upon differentiation (ENRI) (Fig. 1h). A recent study identified two major β-cell subtypes based on H3K27 trimethylation status, with high H3K27me3 β-cells enriched in T2D patients^13^. Using genes upregulated in this subtype (*GCG, ARX, PPY, BUB1, SGCE, SST, ADAMTS18, CPA2, RCN3, CD24*), we defined a high H3K27me3 β-cell population and calculated corresponding module scores. Notably, *SCIN* expression was reduced in this subpopulation as it was expressed in mature β-cells. (Fig. 1i).

Together, these data define SCIN as a villus/L-cell and β-cell–enriched, late-onset factor under Polycomb control, with intestine and β-cell enhancer occupancy by lineage-defining transcription factors, positioning SCIN at the intersection of endocrine identity and glucose-regulated hormone secretion (Fig. 1; Extended Data Figs. 1–3).

### Scinderin acts as a glucose- and PI(4)P-regulated actin modulator at the Golgi Apparatus

Next, we analyzed the intracellular localization of Scinderin. SCIN concentrated at the Golgi in both GLP-1–secreting L-cells and human β-cells: SCIN partially overlapped with GLP-1/proinsulin and showed strong colocalization with the cis-Golgi marker GM130, while showing little overlap with the ER marker calreticulin; fractionation placed SCIN at the ER–Golgi interface (Fig. 2a–g; Extended Data Fig. 4a-e). Consistent with a Golgi-proximal role in actin remodeling, in contrast to the role of the closely related protein Gelsolin (GSN) in cortical actin remodeling, SCIN aligned with perinuclear F-actin patches but not cortical actin in L-cells and β-cells (Fig. 2c, f; Extended Data Fig. 4f).

**Figure 2.**
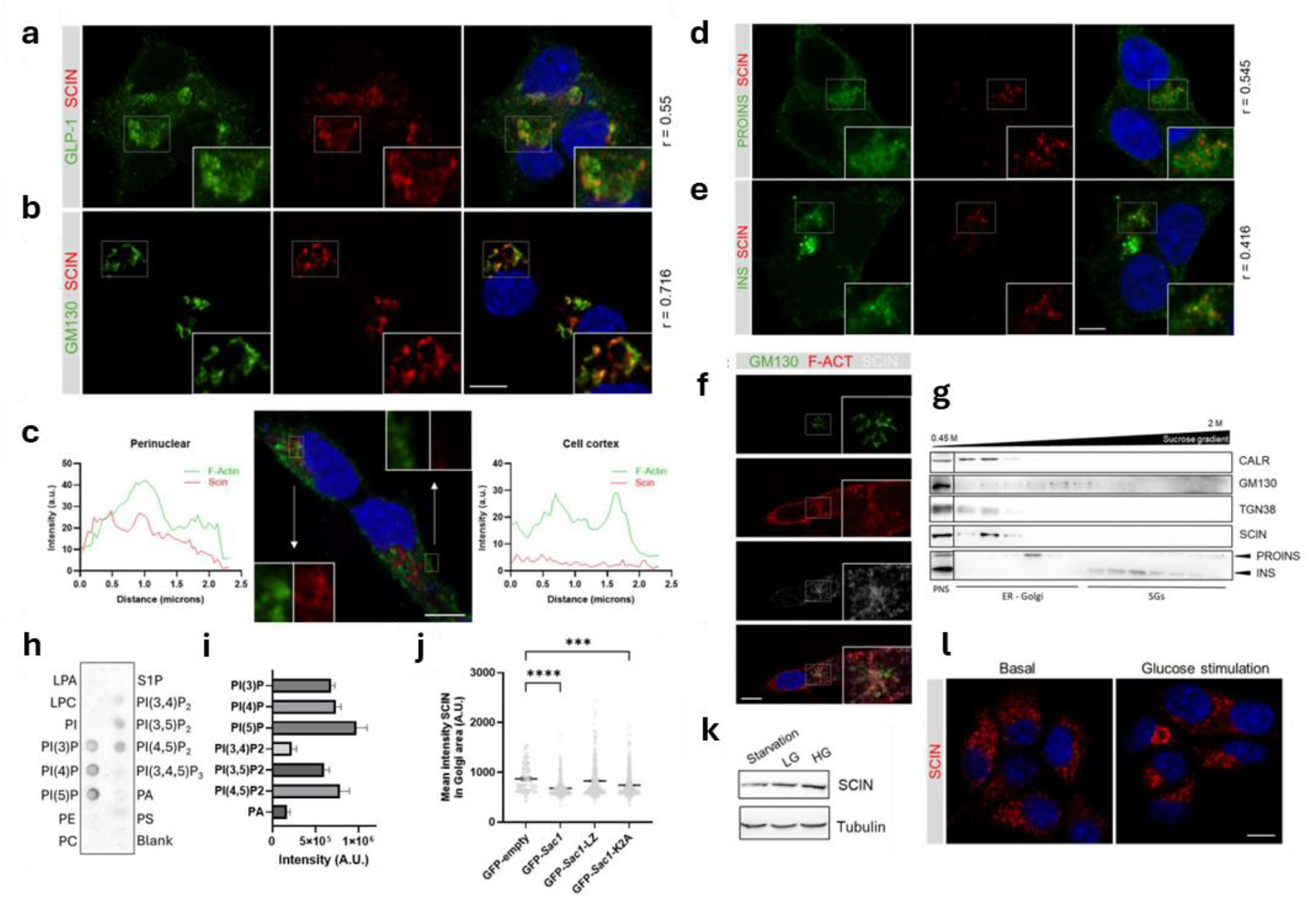
Scinderin is a glucose and PI(4)P-dependent actin remodeller localized to the Golgi. **a–c**, Subcellular localization of Scinderin (SCIN) in enteroendocrine GLUTag cells. **a**, SCIN partially colocalizes with GLP-1. Pearson’s correlation values (n = 5) are shown. **b**, SCIN strongly colocalizes with the Golgi marker GM130. **c**, SCIN overlaps with perinuclear F-actin but not cortical F-actin, as shown by intensity profiles. Scale bar, 5 µm. **d–f**, SCIN localization in human EndoC-βH1 cells. **d, e**, SCIN partially colocalizes with proinsulin and insulin (Pearson’s r, n = 5). **f**, SCIN colocalizes with Golgi-associated F-actin patches (GM130). Scale bars: 5 µm (d,e); 10 µm (f). **g**, Subcellular fractionation of EndoC-βH1 cells. SCIN is enriched at ER–Golgi interface fractions marked by CALR (ER), GM130/TGN38 (Golgi), and PROINS/INS (secretory granules). PNS, post-nuclear supernatant. **h, i**, Lipid-binding assays showing SCIN preferential interaction with phosphatidylinositol phosphates (PIPs) (n = 2). **j**, PI(4)P depletion modifies SCIN Golgi localization. GLUTag cells expressing WT SAC1 or Golgi-localized (K2A) but not ER- (LZ) SAC1 variants showed reduced SCIN Golgi intensity (n = 380–500). Data are mean ± SEM; ***p ≤ 0.001, ****p ≤ 0.0001 (Kruskal–Wallis with Dunn’s post-hoc test). **k, l**, Glucose regulates SCIN protein levels and morphology. **k**, Immunoblot of EndoC-βH1 cells cultured in 0, 5.6, or 20 mM glucose for 4 h shows glucose-dependent SCIN upregulation. **l**, MIN6 cells show punctate SCIN under basal glucose (3 mM) and elongated SCIN structures after glucose stimulation (20 mM). Scale bar, 10 µm.

When we subjected recombinant SCIN to lipid dot-blot we found that it bound phosphoinositide lipids, with a clear preference for Golgi-enriched PIPs including PI(4)P (Fig. 2h, i; Extended Data Fig. 5a). Consistent to this, perturbing PI(4)P confirmed lipid-dependent targeting as overexpression of SAC1 (WT) or a Golgi-restricted SAC1(K2A) depleted Golgi PI(4)P and reduced SCIN signal in the Golgi region, whereas ER-restricted SAC1(LZ) had minimal effect (Fig. 2j; Extended Data Fig. 5b).

Glucose acutely regulated SCIN abundance and organization as high glucose increased SCIN protein in EndoC-βH1 cells and converted punctate SCIN into elongated Golgi-associated structures in MIN6 cells (Fig. 2k, l). Blocking anterograde traffic with brefeldin A trapped SCIN-positive vesicles at the Golgi, indicating that SCIN-decorated carriers bud from this organelle (Extended Data Fig. 5c).

We also noticed that at the transcript level, *SCIN* mRNA was relatively stable in β-cells and rose with glucose, whereas Golgi stress downregulated *SCIN* in human islets (Extended Data Fig. 5d– f).

Together, these data identify SCIN as a glucose-responsive, PI(4)P-directed actin modulator that operates at the Golgi/ER–Golgi interface to coordinate Golgi-associated actin and vesicle biogenesis in endocrine cells.

### Scinderin is required for Golgi structural organization and prohormone trafficking

Because our results show that SCIN operates at the ER–Golgi interface, we next evaluated whether its depletion perturbs Golgi integrity and the flow of cargo through the early secretory pathway. In GLUTag L-cells, SCIN deletion caused retention of GLP-1/ProGCG within the Golgi, with enlarged GLP-1–positive Golgi regions and increased signal intensity; concomitantly, GM130-positive Golgi elements expanded, while ERGIC-53 structures were reduced and more compact, consistent with impaired ER-to-Golgi flux (Fig. 3a–c). These phenotypes concur with the broader transcriptional response of SCIN-deficient GLUTag cells, in which DEGs and GO terms converge on membrane trafficking and organelle pathways (Extended Data Fig. 6).

**Figure 3.**
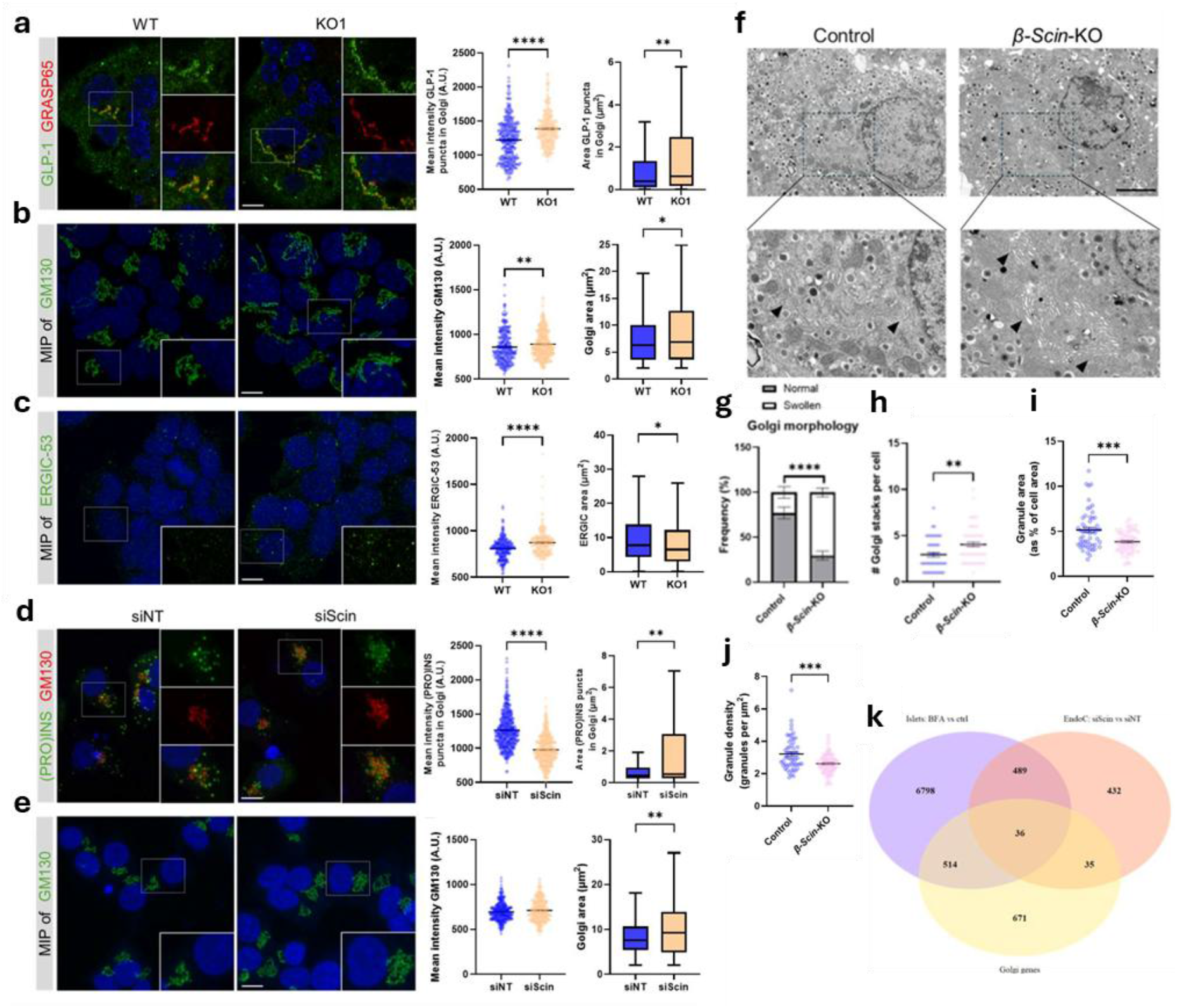
SCIN maintains Golgi architecture and prohormone trafficking. **a–c**, Immunofluorescence of WT and *Scin* KO1 GLUTag cells stained for GLP-1 with GRASP65 (a), GM130 (b), or ERGIC-53 (c). *Scin* loss caused GLP-1 retention in the Golgi, enlarged GM130-positive Golgi, and reduced ERGIC-53 structures. Quantification of puncta intensity and area is shown (n = 227–393). **d–e**, EndoC-βH1 cells transfected with control or *SCIN* siRNA stained for (Pro)insulin with GM130 (d) or GM130 alone (e). SCIN knockdown increased Golgi-associated (Pro)insulin area but reduced intensity, and expanded the Golgi (n = 299–435). **f–j**, Electron micrographs of β-cell Golgi in control and *β-Scin-KO* mice (f). *β-Scin-KO* cells showed more swollen Golgi (g), increased Golgi stack number (h), and reduced dense-core granule area and number (i,j) (n = 55–60 cells). Scale bars, 5 µm (a–e), 2 µm (f). Data are mean ± SEM; *p* ≤ 0.05, **p** ≤ 0.01, \***p** ≤ 0.001, \*\***p** ≤ 0.0001 (Mann–Whitney U test; χ^2^ for g). **k**, Differentially expressed genes (DEGs) from siScin-treated EndoC-βH1 cells were intersected with DEGs from brefeldin-A–treated islets and Golgi-associated genes, identifying 525 shared DEGs, including 36 common Golgi-associated genes; 35 additional Golgi-associated DEGs were unique to SCIN knockdown, yielding 71 Golgi-associated DEGs in total.

In human β-cells, siRNA-mediated *SCIN* knockdown (siScin) increased the Golgi-associated (Pro)insulin area but reduced its intensity and expanded GM130-positive membranes, indicating a trafficking bottleneck at the Golgi (Fig. 3d–e). Transcriptome profiling of siScin EndoC-βH1 cells revealed deregulation of organelle and trafficking programs (Extended Data Fig. 7), and intersection of these DEGs with brefeldin-A (BFA)–treated islets^17^ and a curated Golgi gene set identified 71 Golgi-associated DEGs, including 36 shared with the BFA response (Fig. 3k; Extended Data Fig. 8a), supporting a Golgi-stress–like state upon SCIN loss. *SCIN*-silenced EndoC-βH1 cells were also more sensitive to low-dose BFA, functionally linking SCIN deficiency to compromised Golgi function (Extended Data Fig. 8b).

*In vivo, β-Scin-KO* islets exhibited swollen and disorganized Golgi with an increased number of stacks per cell and reduced dense-core granule area and number by electron microscopy (Fig. 3f–j). Bulk RNA-seq from *β-Scin-KO* islets reinforced these structural and functional defects by enriching for vesicular trafficking categories (Extended Data Fig. 9). Also, bulk RNA-seq from intestinal *Scin*-KO mouse organoids reveal differentially expressed genes (e.g. CES1B, LPCAT2 and ST3GAL6) involved in ER-Golgi membrane-related processes (Extended Data Fig. 10). Together, these data position SCIN as a PI(4)P-directed, Golgi–associated actin remodeler essential for maintaining Golgi architecture, sustaining ER–Golgi trafficking, and supporting prohormone export and granule biogenesis in endocrine cells.

### Scinderin controls hormone content, Golgi F-actin remodelling, and nutrient-stimulated secretion

Loss of SCIN impaired endocrine (pro)hormone stores and nutrient-triggered secretion in both L-cells and β-cells. In GLUTag L-cells, CRISPR-mediated *Scin* deletion reduced ProGCG and intracellular GLP-1 levels (Fig. 4a–c), and KO cells failed to enhance GLP-1 secretion in response to glucose while maintaining normal cAMP-stimulated secretion (Fig. 4d). Interestingly, organoids derived from *Vil-Scin*-KO ileum displayed a significantly reduced glutamine-evoked GLP-1 secretion (Extended Data Fig. 15).

**Figure 4.**
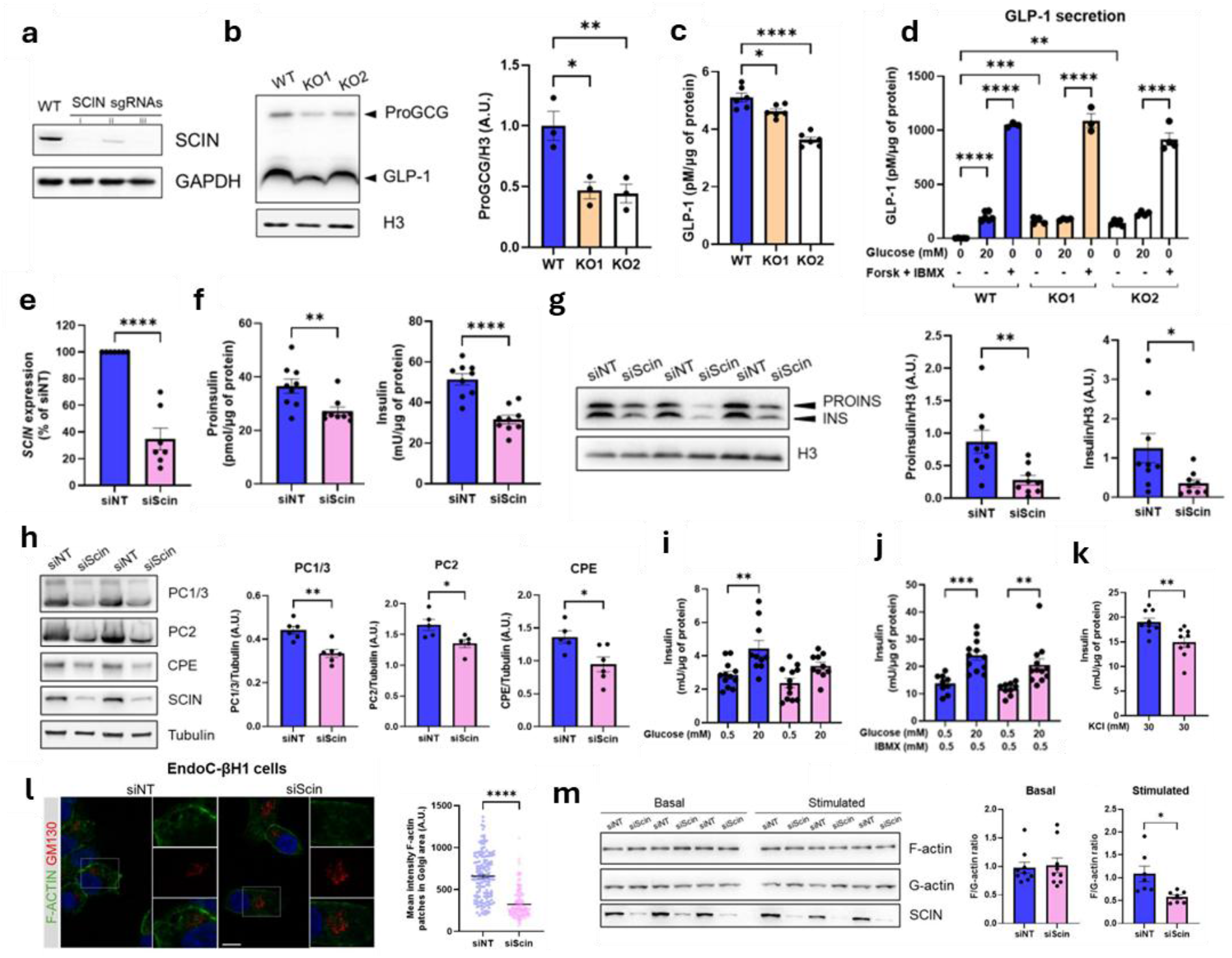
SCIN is required for prohormone content, Golgi F-actin remodelling, and nutrient-triggered secretion. **a–d**, SCIN deletion in GLUTag cells reduces proglucagon/GLP-1 content and abolishes glucose-stimulated secretion. **a**, CRISPR/Cas9 deletion of *Scin* using multiple sgRNAs; sgRNAs i and iii were used as KO1 and KO2. **b–c**, Immunoblot and ELISA quantification of intracellular ProGCG and GLP-1, respectively, show reduced hormone content in KO cells (n = 6). **d**, GLP-1 release under basal (0 mM), glucose-stimulated (20 mM), or forskolin+IBMX conditions. KO cells failed to increase secretion with glucose but retained cAMP responsiveness (n = 3–6). **e–h**, SCIN knockdown in EndoC-βH1 cells reduces (Pro)insulin content and processing enzymes. **e**, Knockdown efficiency by qPCR. **f–g**, ELISA and immunoblot analyses show decreased ProINS and INS levels in siScin cells (n = 9). **h**, Proinsulin-processing enzymes PC1/3, PC2, and CPE are reduced upon SCIN silencing (n = 5– 6). **i–k**, SCIN is required for nutrient-stimulated insulin secretion. **i**, Glucose-stimulated (20 mM) insulin secretion is abolished in SCIN-KD cells (n = 8–12). **j**, IBMX partially restores glucose responsiveness (n = 9–12). **k**, KCl-evoked secretion is diminished in siScin cells. **l–m**, SCIN regulates Golgi-associated F-actin remodelling. **l**, F-actin and GM130 staining in EndoC-βH1 cells shows altered Golgi F-actin intensity in siScin cells (n = 137–160). Scale bar, 5 μm. **m**, Biochemical F/G-actin fractionation reveals normal basal actin ratios but defective stimulus-induced F-actin remodelling in SCIN-KD cells (n = 7–9). Data are mean ± SEM; \**p* ≤ 0.05, **p ≤ 0.01, ***p ≤ 0.001, ****p ≤ 0.0001 by appropriate statistical tests: one-way ANOVA with Dunnet’s post-hoc test (c), with Šidák’s post-hoc test (d, i, j); Unpaired two-tailed t-test (e-f, h, k); Mann-Whitney U test (g, l-m).

*SCIN* silencing in human EndoC-βH1 cells decreased (Pro)insulin content and concomitantly prohormone-processing enzymes PC1/3, PC2, and CPE (Fig. 4e–h), consistent with the aforementioned vesicular trafficking defect (Fig. 3d, i-j). A similar defect was observed *ex vivo*: *β-Scin-KO* islets exhibited normal morphology but markedly reduced (pro)insulin content by immunoblot and ELISA (Extended Data Fig. 11a–c). Human EndoC-βH1 cells where *SCIN* was silenced exhibited an impaired glucose-stimulated insulin secretion (GSIS) response, while IBMX restored secretory competence, indicating that the glucose-triggering pathway, rather than the cAMP-amplifying pathway, was selectively compromised (Fig. 4i–j). Importantly, SCIN loss blunted depolarization-evoked secretion, with siScin cells releasing markedly less insulin upon KCl stimulation (Fig. 4k), indicating impaired calcium-dependent triggering downstream of glucose sensing. MIN6 β-cells showed the same phenotype: reduced insulin content and absent GSIS following SCIN depletion, with IBMX restoring its responsiveness (Extended Data Fig. 13), demonstrating that SCIN function is conserved across species and β-cell models.

Mechanistically, SCIN loss altered actin remodelling at the Golgi. In EndoC-βH1 cells, SCIN knockdown decreased Golgi-associated F-actin intensity and impaired stimulus-induced F/G-actin transitions (Fig. 4l–m). Consistent results were obtained with *Scin*-KO GLUTag cells where cells exhibited reduced F-actin accumulation specifically at the Golgi (Extended Data Fig. 12).

Together, these results demonstrate that SCIN is a conserved, Golgi-associated actin remodeler required for producing a correct prohormone output and coupling nutrient stimulation to regulated secretion in both L-cells and β-cells.

### SCIN coordinates enteroendocrine output and β-cell secretion to sustain systemic glucose control in vivo

Tissue-specific SCIN deletion was validated by the absence of protein and locus excision in conditional SCIN-KO models (Extended Data Fig. 14). Vil-Scin-KO mice displayed broadly intact intestinal architecture, with the notable exception of shorter ileal villi and elongated microvilli (Figs. S19–20). Their body weight and composition were largely unchanged (Extended Data Fig. 16), and basal food and water intake was comparable (Extended Data Fig. 18). *In vivo* intestinal SCIN loss blunts the incretin response and glucose tolerance: during oGTT, *Vil-Scin*-KO mice of both genders exhibited higher glycaemia—most prominently at 15–30 min—and failed to raise plasma GLP 1 (Fig. 5a–d), with parallel defects in PYY (Extended Data Fig. 17) despite preserved insulin response, islet morphology, normal insulin sensitivity and gastric emptying (Extended Data Figs. 24a-c, 27).

**Figure 5.**
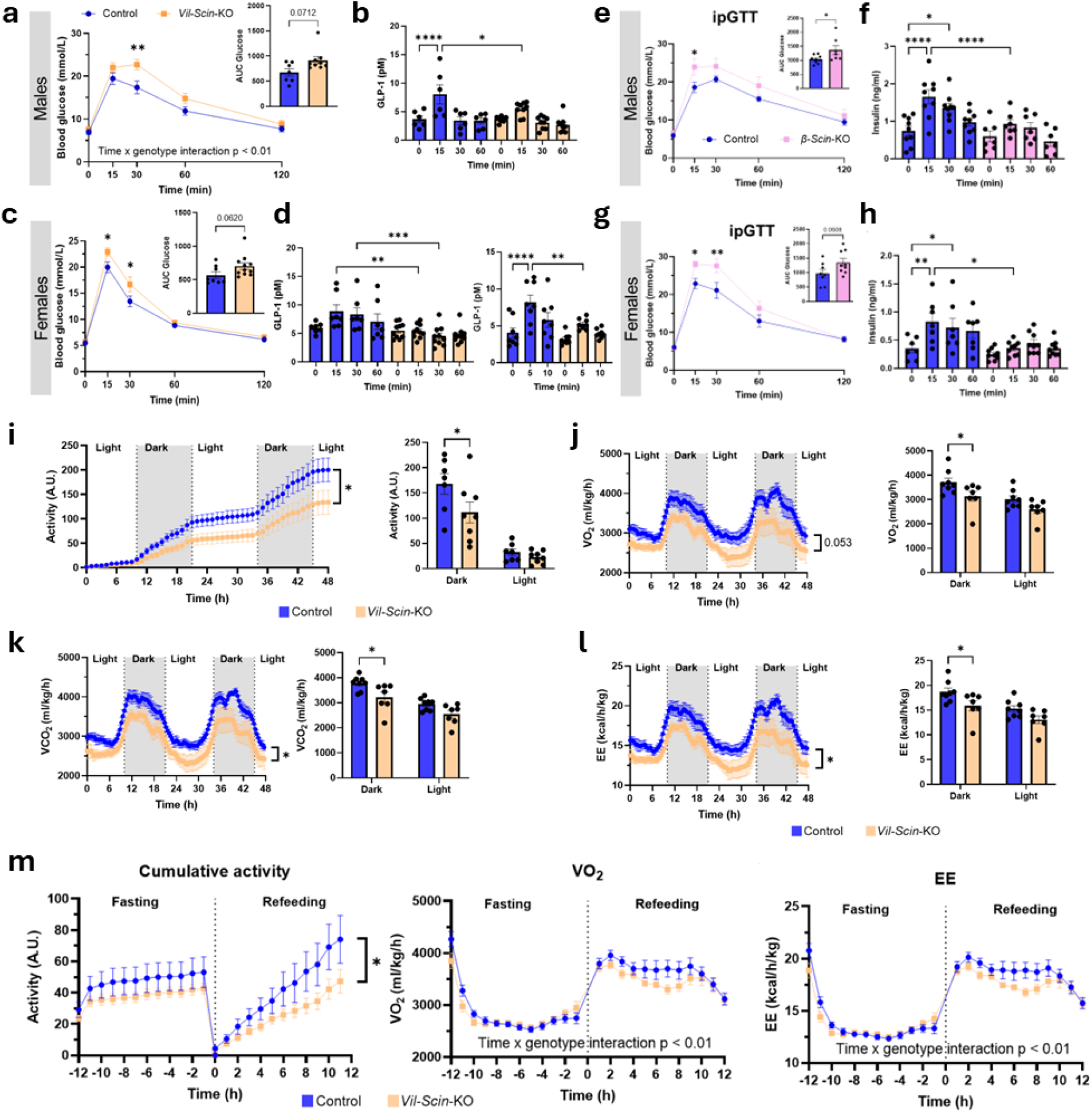
Scinderin coordinates enteroendocrine and pancreatic endocrine responses in vivo. **a–d**, Glucose tolerance and incretin responses in *Vil-Scin-KO* mice. **a, c**, Male (a) and female (c) mice subjected to oral glucose tolerance tests (oGTT) displayed impaired glucose clearance, with elevated glycemia at 15–30 min (n = 8–12). **b, d**, Plasma GLP-1 failed to rise after glucose in both sexes of *Vil-Scin-KO* mice (n = 6–9). **e–h**, Glycemic and insulin responses in *β-Scin-KO* mice. **e, g**, Male (e) and female (g) *β-Scin-KO* mice exhibited elevated blood glucose during intraperitoneal glucose tolerance tests (ipGTT) (n = 7–9). f,h, Insulin secretion was abolished in *β-Scin-KO* mice during ipGTT (n = 7–9). **i–**l, Activity and metabolic rate in *Vil-Scin-KO* mice. **i**, Cumulative activity over 48 h revealed reduced locomotion, especially during the dark phase (n = 7–8). **j–l**, Indirect calorimetry showed decreased oxygen consumption (VO_2_) (j), carbon dioxide production (VCO_2_) (k), and energy expenditure (EE) (l), most prominently during the dark phase (n = 7–8). **m**, Fasting–refeeding responses. Locomotor activity diverged between genotypes upon refeeding (n = 8). VO_2_ and EE exhibited altered postprandial dynamics in *Vil-Scin-KO* mice, indicating defective nutrient-driven metabolic activation. Data are mean ± SEM; statistical tests as indicated in original figure panels. Data are presented as mean ± SEM. *p ≤ 0.05, **p ≤ 0.01, ***p ≤ 0.001, ****p ≤ 0.0001 vs. indicated groups by two-way repeated-measures ANOVA with Šidák’s test (A, C, E, G), Mann-Whitney U test (AUC of A, C), unpaired t test (AUC of E, G), one-way ANOVA repeated-measures with Šidák’s post-hoc test (B, D, F, H) and by two-way ANOVA repeated-measures with Šidák’s post-hoc test (i-m)

In the pancreas, β-cell SCIN deletion impaired glucose handling and stimulus–secretion coupling: *β-Scin-KO* mice showed elevated glycaemia during ipGTT (but not oGTT; Extended Data Fig. 23) and failed to augment plasma insulin (Fig. 5e–h). Additionally, *β-Scin-KO* displayed reduced islet mass and cell number but preserved α/β areas (Figs. S24d-h, S25, S29), slightly lower body weight and normal tissue weights (Extended Data Fig. 21). *β-Scin-KO* mice showed unchanged *ad libitum* glycaemia, and a trend toward greater insulin sensitivity in ITT whereas food and water intake was similar (Extended Data Figs. 22, 30).

At the whole-body level, SCIN deficiency in the intestine reduced postprandial activity and energy expenditure. In metabolic cages, *Vil-Scin-KO* mice displayed lower locomotor activity (48 h; dark-phase predominant; Fig. 5i) alongside decreased VO_2_, VCO_2_, and EE (Fig. 5j–l), with similar food/water intake (Extended Data Fig. 18). Upon a fasting–refeeding experiment, phenotypic differences emerged at refeeding: *Vil-Scin-KO* mice showed blunted rises in activity, VO_2_, and EE (Fig. 5m–n), consistent with altered postprandial metabolic activation; body-temperature mapping showed parallel, non-significant downward trends (Extended Data Fig. 26). *Vil-Scin-KO* showed a trend toward reduced intestinal GLP-1 content without changes in L-cell number, indicating decreased hormone content per cell, consistent with our *in vitro* findings (Extended Data Fig. 28 c-g). Additional intestinal analyses supported a selective enteroendocrine defect, as gut macroscopic parameters were largely preserved. (Extended Data Fig. 28 a, b). Moreover, as a previous study suggested that IELs can influence energy expenditure by modulating GLP-1 bioavailability^18^, concept later challenged^19^, our flow-cytometric analyses revealed no alterations in intestinal lymphocyte populations, indicating that the metabolic phenotype does not arise from immune cell changes (Extended Data Fig. 31).

Together these *in vivo* results establish SCIN as a unifying regulator that couples enteroendocrine GLP-1 output to β-cell insulin secretion and thereby maintains postprandial glucose control and energy expenditure.

### Scinderin aligns with diabetes-associated β-cell state shifts

To connect our mechanistic findings with human disease, we analyzed multiple human β-cell transcriptomic datasets^16,20^. Bulk RNA-seq of islets from individuals with type 1 diabetes (T1D) and pseudobulk analysis of beta-cell clusters from scRNA-seq data in type 2 diabetes (T2D) reveal reduced SCIN expression, accompanied by a parallel decrease in INS in both conditions, consistent with a shared β-cell dysfunctional state that includes impaired secretory granule maturation (Fig. 6a-b). Furthermore, genes upregulated upon *SCIN* loss in human β-cells were significantly enriched among genes elevated in diabetes-associated, low-maturity β-cell clusters (Fig. 6c), including those enriched for high H3K27me3 signatures and SCIN-associated Golgi stress genes (Fig. 6d). Interestingly, we observed that Golgi apparatus (GA) genes upregulated upon BFA treatment in islets (padj < 0.01 and log2FC > 4), along with canonical ER stress markers (DIT3, HSPA5, ATF4, and XBP1), were enriched in cluster 8 (Extended Data Fig. 32). This enrichment appeared independent of H3K27 trimethylation status and mature Beta cell identity, suggesting that not all ER stress responses are associated with SCIN expression or PRC2 activity. Together, these analyses position loss of the PRC2 target SCIN as a key axis underlying conserved β-cell dysfunction in human diabetes, encompassing Golgi stress, loss of maturation, and diminished secretory competence.

**Figure 6.**
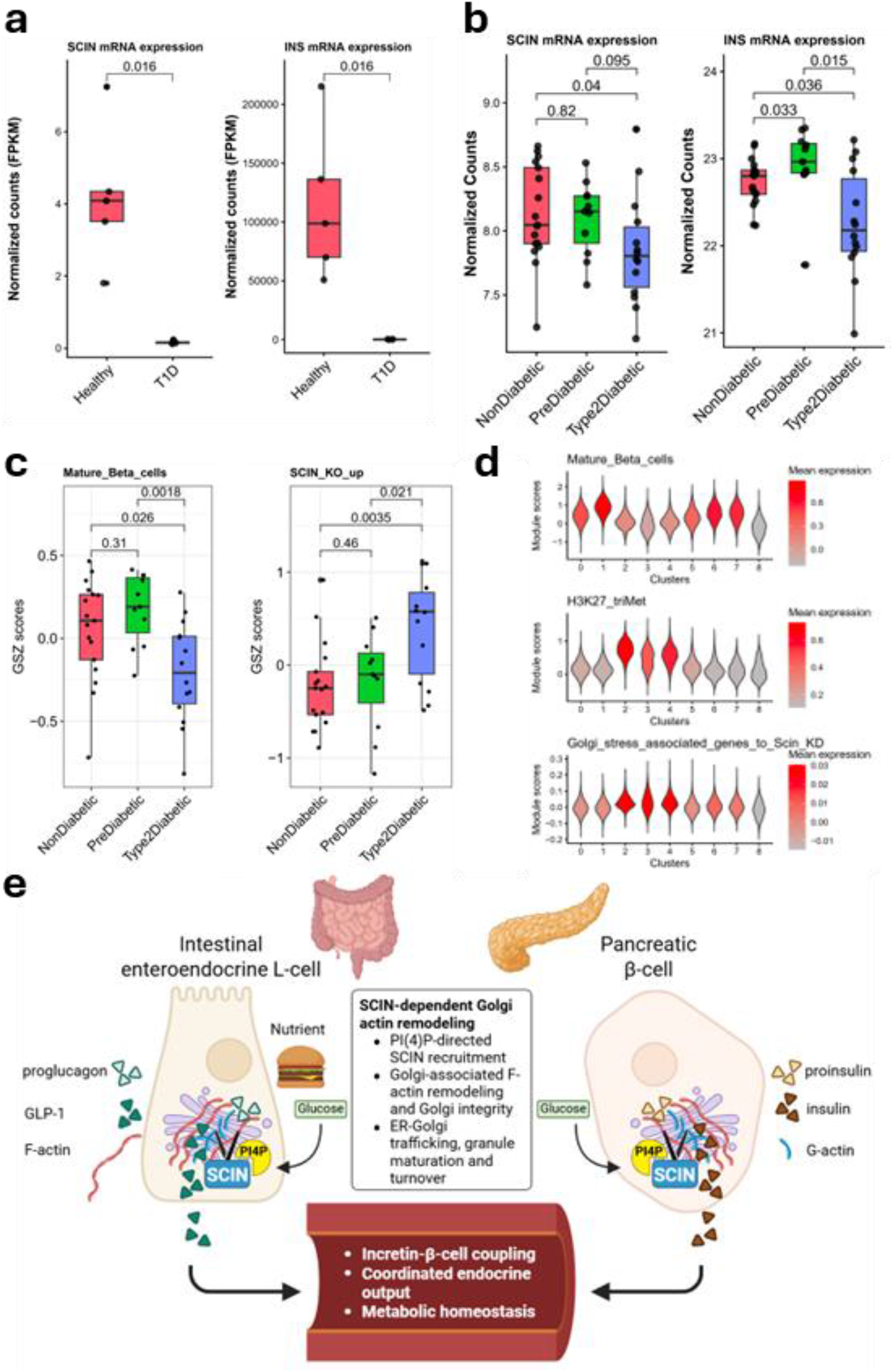
SCIN expression tracks mature-to-stress β-cell state shifts in human diabetes. **a**, Bulk RNA-seq data from healthy and T1D islets show reduced SCIN and INS expression. **b**, Pseudobulk analysis of the beta-cell cluster from scRNA-seq data also shows reduced SCIN and INS expression in T2D donors compared with non-diabetic donors. Data were normalized using FPKM (a) and DESeq2 VST (b). Statistical significance was assessed using the Mann-Whitney test with Benjamini-Hochberg (BH) correction for multiple comparisons. Points represent individual donors, with median ± interquartile range (IQR) indicated. **c**, Gene Set Z-score (GSZ) analysis shows that top 20 genes upregulated upon *SCIN* loss in EndoC-βH1 cells are significantly enriched among genes elevated in T2D β-cell clusters characterized by reduced maturity and increased stress. **d**, β-cell clusters 2, 3, and 4 (Extended Data Fig. 32) enriched for high H3K27me3 signatures and low β-cell maturity, also showed enrichment for Golgi apparatus (GA)-associated stress genes shared with siScin EndoC-βH1-genes in Fig. 3k differentially expressed in SCIN knockout (KO) conditions. **e**, Model of SCIN-dependent Golgi actin remodeling in endocrine secretory control. SCIN localizes to PI(4)P-enriched Golgi membranes in intestinal enteroendocrine L-cells and pancreatic β-cells, where it regulates Golgi-associated F-actin remodeling and sustains Golgi integrity. This actin-dependent organization supports efficient ER– Golgi trafficking, prohormone processing, granule maturation, and the generation of stimulus-competent secretory granules. In L-cells, SCIN-dependent Golgi actin remodeling enables nutrient-evoked GLP-1 secretion, while in β-cells it sustains glucose-triggered insulin release. Coordinated enteroendocrine and pancreatic endocrine output integrates incretin–β-cell coupling to ensure postprandial glucose control and systemic metabolic homeostasis

## Discussion

Efficient metabolic control requires endocrine cells to rapidly translate nutrient availability into hormone biosynthesis and secretion. In both intestinal enteroendocrine cells and pancreatic β-cells, this process relies not only on stimulus–exocytosis coupling at the plasma membrane but also on the structural and functional integrity of the Golgi apparatus and its capacity to generate secretion-competent granules. Although actin dynamics have long been implicated in regulated secretion, previous models have largely emphasized cortical actin remodeling during granule docking and fusion^6,8^. Our findings position Scinderin (SCIN) as a lineage-restricted actin remodeller that operates earlier in the secretory pathway, at the Golgi–actin interface, to sustain regulated hormone secretion across both intestinal L-cells and pancreatic β-cells.

SCIN localizes to PI(4)P-enriched Golgi membranes and associated F-actin patches, distinct from plasma-membrane actin structures described for gelsolin, myosins, and SNARE-associated regulators^21,22^. This localization places SCIN within a growing class of actin regulators required for Golgi structure and vesicle biogenesis, including MYO1B, MYO18A, and the GOLPH3 complex^23– 25^. Perturbation of these proteins produces Golgi distension, delayed cargo export, and reduced secretory granule formation—phenotypes strikingly similar to those observed upon SCIN loss. Our data therefore extend prior work by identifying SCIN as a dynamic actin severing and nucleating factor that enables Golgi-associated actin turnover required for efficient hormone trafficking.

SCIN recruitment to the Golgi is nutritionally regulated and depends on PI(4)P, a lipid that coordinates actin dynamics and vesicle formation at the trans-Golgi network^26,27^. PI(4)P depletion—either through glucose withdrawal or SAC1 phosphatase expression—displaces SCIN from the Golgi, supporting a model in which nutrient availability modulates Golgi mechanics through lipid-directed actin remodeling. This mechanism aligns with recent findings that PI(4)P-regulating proteins such as PITPNA are reduced in islets from individuals with type 2 diabetes, leading to impaired granule biogenesis and defective glucose-stimulated insulin secretion^28^. Together, these observations suggest that PI(4)P-dependent actin remodeling constitutes a critical regulatory nexus linking nutrient sensing to secretory competence.

Across both L-cells and β-cells, SCIN deficiency causes Golgi swelling, reduced ER-Golgi intermediate compartment structures, retention of prohormones such as ProGCG and proinsulin within the Golgi, and a marked loss of dense-core secretory granules. Similar Golgi alterations have been reported in β-cells under metabolic stress conditions, including high-fat diet feeding, cytokine exposure, and diabetes, where delayed proinsulin trafficking correlates with impaired secretory output^17,29,30^. Transcriptomic analysis further reveals that SCIN loss elicits a conserved Golgi stress response overlapping substantially with brefeldin A–treated islets, without activating canonical ER stress pathways^17,31^. These findings argue that SCIN depletion induces a Golgi-intrinsic stress state, likely driven by defective actin-dependent vesicle budding rather than protein misfolding in the ER.

Functionally, SCIN is indispensable for nutrient-stimulated hormone secretion. In both L-cells and β-cells, loss of SCIN selectively impairs glucose-evoked GLP-1 and insulin release, while cAMP-dependent secretion remains intact. This dissociation mirrors earlier observations in cells lacking actin regulators such as gelsolin or RICTOR, where stimulus-dependent secretion is compromised but can be rescued by cAMP signaling ^21,32^. These findings indicate that SCIN acts upstream of exocytosis, ensuring the availability and turnover of secretion-competent granules rather than directly controlling fusion machinery.

In β-cells, the selective loss of glucose-stimulated insulin secretion likely reflects diminished granule supply. Newly generated granules preferentially support glucose-stimulated insulin secretion and contribute disproportionately to newcomer-granule fusion events^33–35^. Delayed Golgi export and reduced granule density observed upon SCIN depletion would therefore be expected to impair this mode of secretion while sparing cAMP-amplified pathways that enhance recruitment and fusion probability^36,37^. Similar principles likely apply in L-cells, where actin remodeling governs time-dependent granule fusion competence during GLP-1 secretion^7^.

*In vivo*, SCIN deletion reveals its cell-intrinsic importance in endocrine physiology. Intestinal SCIN knockout blunts postprandial GLP-1 and PYY secretion, reduces energy expenditure, and diminishes locomotor activity without altering feeding behavior, phenotypes consistent with impaired incretin and PYY signaling^38–40^. Conversely, β-cell-specific SCIN loss selectively impairs glucose-stimulated insulin secretion while preserving systemic glycemic control through enhanced insulin sensitivity and incretin-mediated compensation, analogous to phenotypes reported in models of actin remodeling and vesicle trafficking defects^32,41^. These observations emphasize that Golgi-actin integrity contributes to endocrine output but can be partially buffered by systemic adaptive mechanisms under physiological conditions.

A defining feature of *SCIN* is its tightly restricted expression pattern. *SCIN* is epigenetically repressed by PRC2 in intestinal crypts and early pancreatic progenitors, and becomes expressed only in terminally differentiated villus L-cells and mature β-cells. This selective activation may reflect shared elements of the mature endocrine transcriptional program. In β-cells, the *SCIN* promoter is bound by core endocrine regulators such as PDX1, NKX2-2 and NKX6-1, while L-cells share overlapping endocrine transcription factors (including NEUROD1, ISL1 and PAX6) that, together with PRC2 release during terminal differentiation, could permit convergent derepression of *SCIN* in both lineages. This developmental gating also distinguishes *SCIN* from ubiquitously expressed actin regulators such as gelsolin^42^ and suggests that *SCIN* integrates endocrine identity with secretory capacity. Consistent with this view, human single-cell datasets reveal reduced *SCIN* expression in stress-associated β-cell states in both type 1 and type 2 diabetes, paralleling the Golgi stress signatures and granule deficits observed experimentally^17^.

While these human data are correlative, their concordance with our genetic and functional perturbations supports a model in which SCIN-dependent Golgi integrity is a key determinant of endocrine secretory fitness. Notably, most current diabetes therapies act downstream, improving insulin sensitivity or amplifying secretion without correcting upstream defects in hormone biosynthesis, granule biogenesis, or Golgi homeostasis^12^. Our findings therefore identify Golgi-associated actin remodeling as an underappreciated vulnerability in metabolic disease and position the SCIN–Golgi axis as a potential target for preserving endocrine cell function.

Several limitations should be acknowledged. Although our data establish PI(4)P-dependent recruitment of SCIN to the Golgi, the molecular partners coupling SCIN to actin filaments and vesicle budding machinery remain undefined. Additionally, while transcriptional and ultrastructural analyses implicate defective actin turnover as a driver of Golgi stress, the causal hierarchy between actin remodeling, lipid organization, and vesicle scission remains unresolved. Finally, compensatory adaptations in murine intestine and islets may attenuate the impact of SCIN loss compared with human disease states. Direct visualization of Golgi-associated actin dynamics and granule trafficking will be essential to refine this model.

In summary, our findings identify SCIN as a conserved, PI(4)P-directed actin remodeller that maintains Golgi organization, prohormone trafficking, and nutrient-evoked hormone secretion in intestinal and pancreatic endocrine cells (Fig. 6e). These results reveal a fundamental requirement for Golgi-associated actin dynamics in metabolic homeostasis and suggest that failure of this machinery may contribute to endocrine fragility in diabetes.

## Supporting information

Additional references

## Resource availability

### Materials availability

All materials used in this paper are available from the lead contact upon request.

### Data and code availability

All the data supporting the findings of this study are available within the article and its supplemental information. Transcriptomic data generated from EndoC-βH1, intestinal organoids and pancreatic islets are deposited in the Gene Expression Omnibus (GEO) database repository: GSE### and are publicly available as of the date of publication. Any additional information required to reanalyze the data reported in this paper is available from the lead contact upon request.

## Acknowledgements

The authors gratefully acknowledge Michael Glogauer (University of Toronto, Canada) for providing the Scin floxed mice, Daniel Drucker (University of Toronto, Canada) for GLUTag cells, Pekka Katajisto (University of Helsinki, Finland) for HEK293 cells, Kari Alitalo (University of Helsinki, Finland) for VilCre mice, and Neale Ridgway (Dalhousie University, Canada) for the SAC1 plasmids. We thank Eini Eskola, Nina Koivisto, Soili Peltomäki, Roza Parveen, Mikko Oittinen, Vikash Chandra, Sarah Baghestani, and Jilab Inc for excellent technical support. The authors acknowledge Biocenter Finland (BF) for services provided through the Tampere Imaging Facility (TIF), the Tampere University Flow Cytometry Facility (TFCF), and Tampere Protein Services. We also acknowledge the Tampere University Histocore and Preclinical Facility for their services. We also thank the Electron Microscopy Unit of the Institute of Biotechnology, University of Helsinki, for providing laboratory facilities. This study was supported by the Research Council of Finland (Grant no. 265575, 370828 and 310011), University of Oulu and Research Council of Finland Profi8 (Grant no. 365202), the Finnish Cultural Foundation, the Diabetes Research Foundation, Jalmari ja Rauha Ahokkaan säätio, and the State funding for university-level health research Tampere University Hospital (Wellbeing services county of Pirkanmaa, projects No. T63474, T66984 and 68084). LMD gratefully acknowledges financial support from the Finnish Cultural Foundation, the Diabetes Research Foundation, the Instrumentarium Science Foundation, the Kyllikki and Uolevi Lehikoinen Foundation, the Finnish National Agency for Education, Tampereen tutkimustyön tukisäätiö, and the Doctoral School of the Faculty of Medicine and Health Technology at Tampere University. DB wants to acknowledge Sigrid Jusélius Foundation, https://ror.org/00ckakm23, @003701165704@, Academy Research Fellowship 2024 Research Council of Finland, 361593 and Excellence Emerging Investigator Grant Novo Nordisk Foundation, NNF24OC0089232. RD acknowledges The Magnus Ehrnrooth foundation. MV-R like to thank Jane and Aatos Erkko Foundation, Research Council of Finland (Grant no. 330896), the European Union’s Horizon 2020 research and innovation program under grant agreement No 101017116, project CoCID (Compact Cell-Imaging Device), and Biocenter Finland’s viral gene transfer platform. DMT acknowledges Research Council of Finland (Grant no. 126161) and Sigrid Juselius Foundation.

## Author contributions

LMD and KV conceived the study. LMD designed the study and generated the figures. Data acquisition was performed by LMD, KV, MP, AG, MO, FTAM, JJG, SH, KE, and JS-V. Data analysis and interpretation were carried out by LMD, KV, MP, CC, VD, MO, RD, and DB. ES, JI, JL, HH, TO, MV-R, and DMT provided resources and assisted with the logistics of data collection. The manuscript was drafted by LMD and KV. KV secured funding and supervised the study. All authors read and approved the final manuscript.

