## Supplementary Figures for "Scinderin-driven Golgi Actin Remodeling coordinates GLP-1 and insulin secretion to regulate glucose homeostasis"

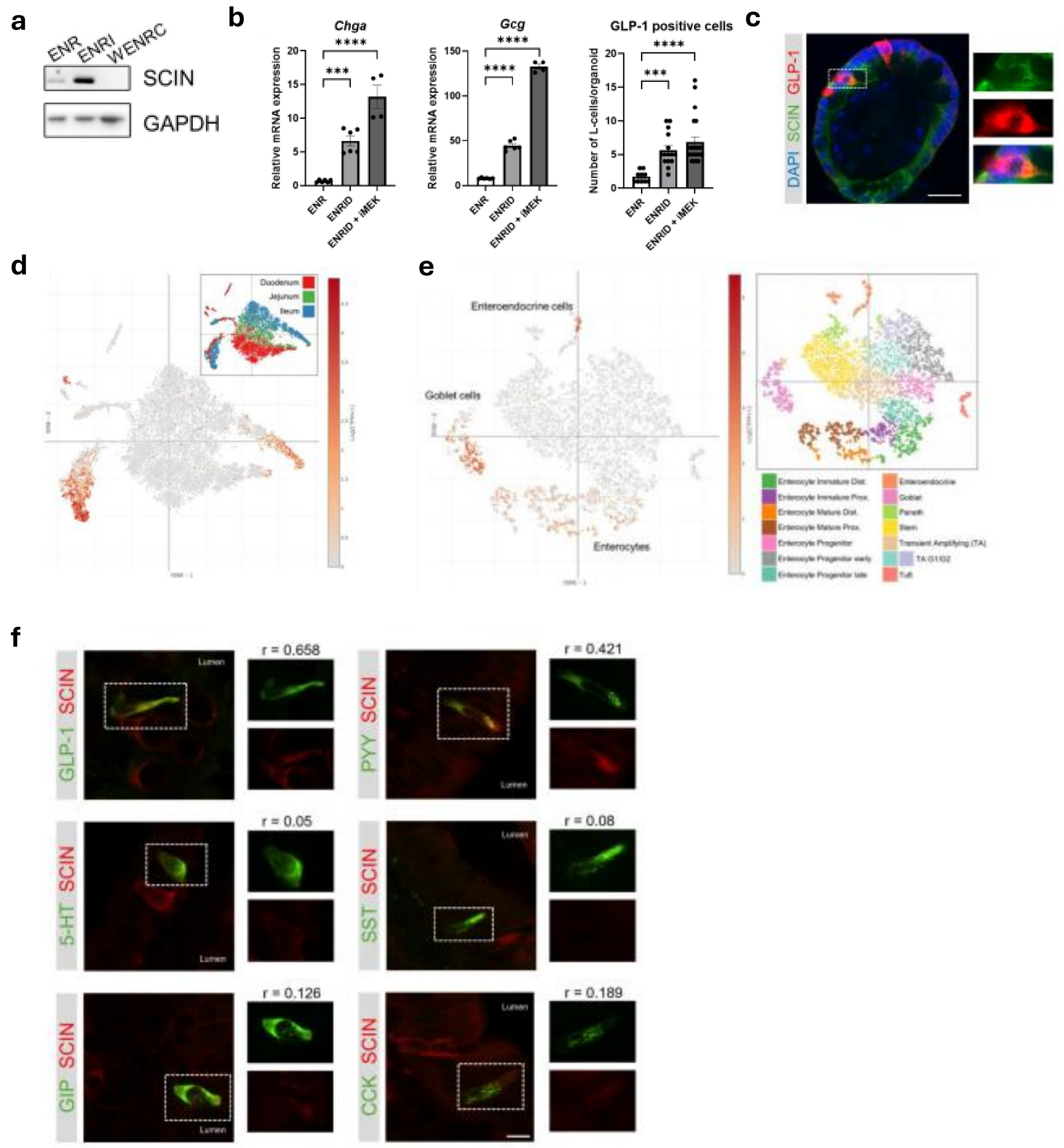


**Extended Data Figure 1. Expression of Scinderin in specific endocrine cells types in the intestine. a,** SCIN protein expression analyzed by immunoblot in mouse intestinal organoids cultured to resemble crypt+villi (ENR), villi (ENRI) and crypt (WENRC). **b,** Evaluation of conditions that enrich for enteroendocrine cells in organoids by supplementing standard ENR medium with Wnt and Notch inhibitors, along with or without EGFR inhibition (referred to as ENRID and ENRID + iMEK, respectively). qPCR analysis of mouse intestinal organoids (n = 4–6) showing expression of the enteroendocrine marker Chromogranin A (Chga, *left*) and L-cell marker Glucagon (Gcg, *middle*). Inhibition of Wnt and Notch signaling significantly increased Chga and Gcg expression, with further enhancement observed upon the addition of EGFR inhibitor. *Right*, organoids were stained for GLP-1 and the number of GLP-1-positive cells per organoid were quantified (n = 10–21). Upon culture in ENRID and ENRID + iMEK media, organoids shown a significant increase in the number of GLP-1-positive cells. Data are presented as mean ± SEM. ***$p \leq 0.001$, ****$p \leq 0.0001$ vs. indicated groups by one-way ANOVA with Dunnett's post-hoc test (qPCR) or Kruskal-Wallis test with Dunn's post-hoc test (GLP-1 staining). **c,** Immunofluorescent staining of mouse intestinal organoids differentiated toward the villus lineage. Scale bar, 20 µm. **d,** Scin expression across mapped intestinal regions of scRNAseq data from mouse small intestine (data from Haber et al., 2017). tSNE plot is used for visualization and the relative expression is shown as log2(TPM+1), where TPM denotes transcripts per million. Top legend, clusters from the tSNE plot colored by intestinal regions. Scin is mainly expressed in the ileum. **e,** tSNE visualization of Scin expression across different intestinal cell types from scRNAseq data of the mouse small intestine (Haber et al., 2017). Data suggest Scin enrichment in enterocytes, goblet cells, and enteroendocrine cells. **f,** Representative immunostaining of SCIN and various enteroendocrine hormones in mouse the intestine. The average Pearson's correlation coefficient from colocalization analysis of different small intestine samples (n = 4) is shown. SCIN colocalizes with GLP-1 and PYY. Scale bar, 10 µm.

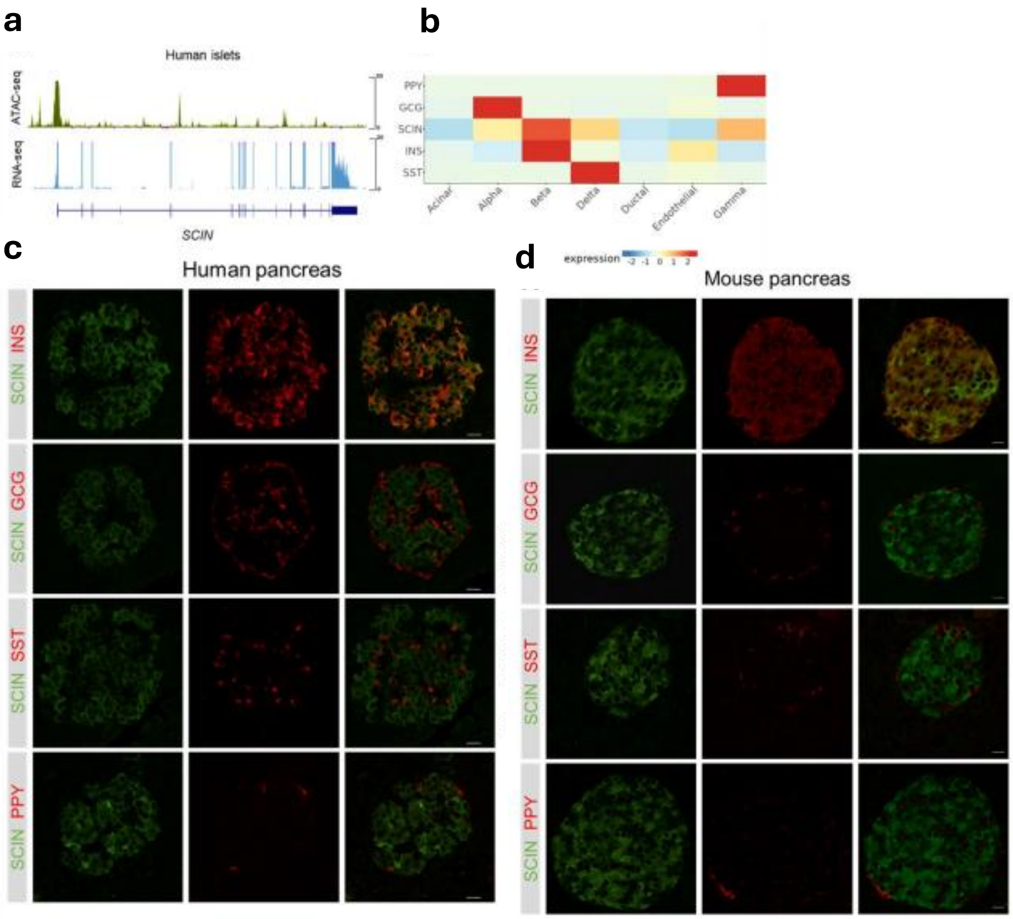


**Extended Data Figure 2**. **Expression of Scinderin in specific endocrine cells types in the pancreas. a,** Normalized SCIN mRNA expression (RNA-seq) and chromatin accessibility (ATAC-seq) at the SCIN locus in human islets (data from Mularoni et al., 2017 – accessed via The Islet Regulome Browser). **b,** Heatmap of SCIN and endocrine hormone mRNA expression across distinct islet cell types: beta (INS), alpha (GCG), delta (SST), and F (PPY) cells. Data from Mummey et al., 2024. **C)** Representative immunofluorescence image of human pancreatic tissue stained with anti-SCIN antibody and hormone-specific antibodies to identify islet cell types. Scale bar, 20 μm.

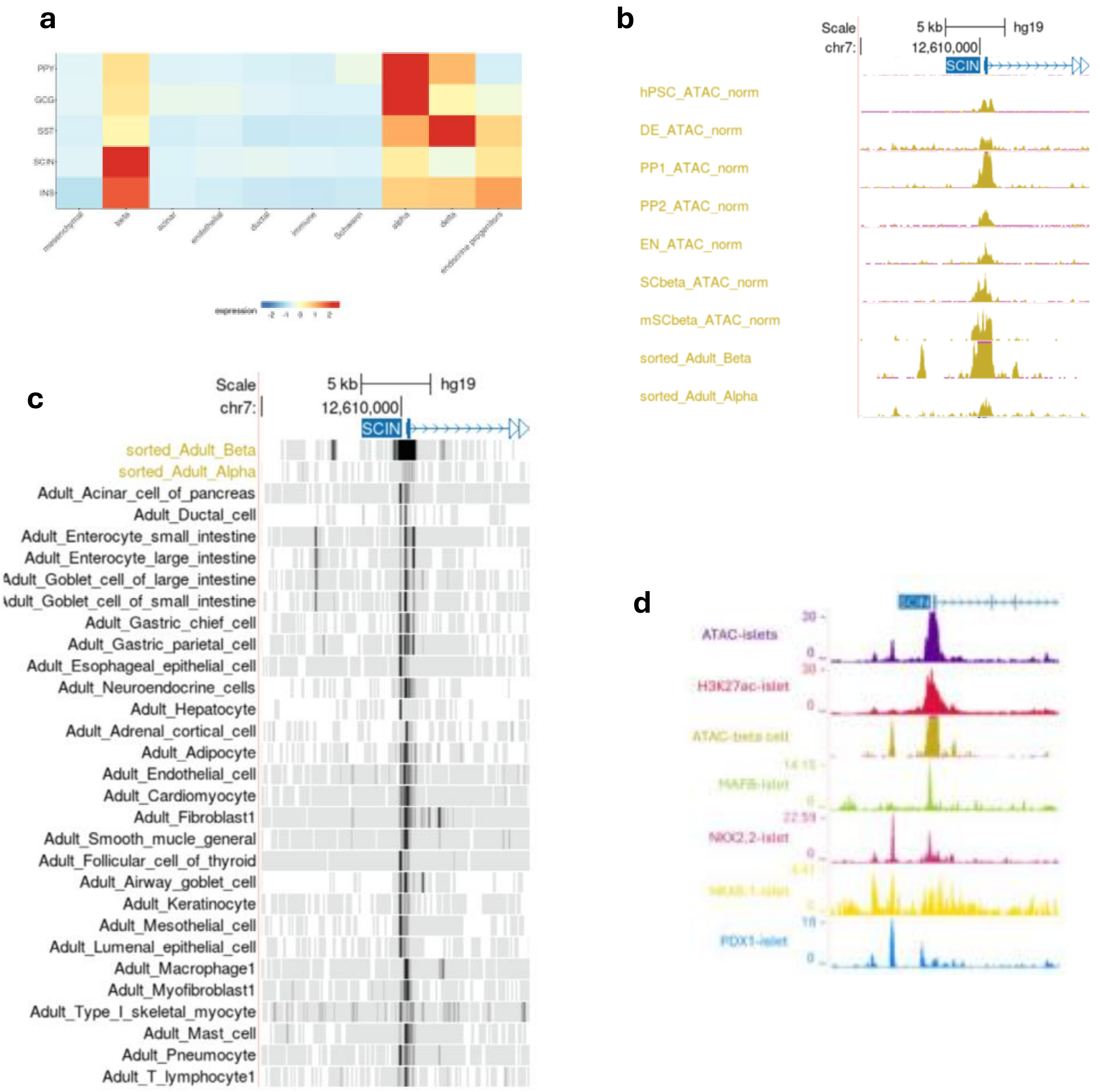


**Extended Data Figure 3**. **Epigenetic regulation of Scinderin expression during human pancreatic development. a,** Heatmap showing SCIN and endocrine hormone mRNA expression across distinct islet cell types in human fetal pancreas at 16 weeks post-conception (data from Olaniru et al., 2023). SCIN expression is enriched in cells corresponding to the fetal pancreatic beta cell lineage at the late endocrine differentiation phase. **b,** Normalized chromatin accessibility (ATAC-seq) on SCIN promoter and upstream genomic region throughout *in vitro* stem cell differentiation into SC-islets (hPSC, DE, PP1, PP2, EN) and in primary beta and alpha cells. Accessibility at the SCIN promoter increases progressively, peaking in sorted primary beta cells. Data from Alvarez-Dominguez et al. 2020. **c,** Chromatin accessibility (bulk and single cell ATAC sequencing) on SCIN promoter and upstream genomic region in sorted primary adult beta and alpha cells compared to other adult tissues. Upstream enhancers are accessible in beta cells and in adult human intestinal cells (enterocyte and goblet cells), whereas these regions are inaccessible in other adult cell types. Single cell ATAC data from Zhang et al 2021. **d,** ATAC-seq profiles for islets and primary beta cells, together with chromatin immunoprecipitation (ChIP-seq) for H3K27ac, MAFB, NKX2-2, NKX6-1, and PDX1. A beta cell specific upstream enhancer at the SCIN locus is bound by NKX2-2, NKX6-1 and PDX1.

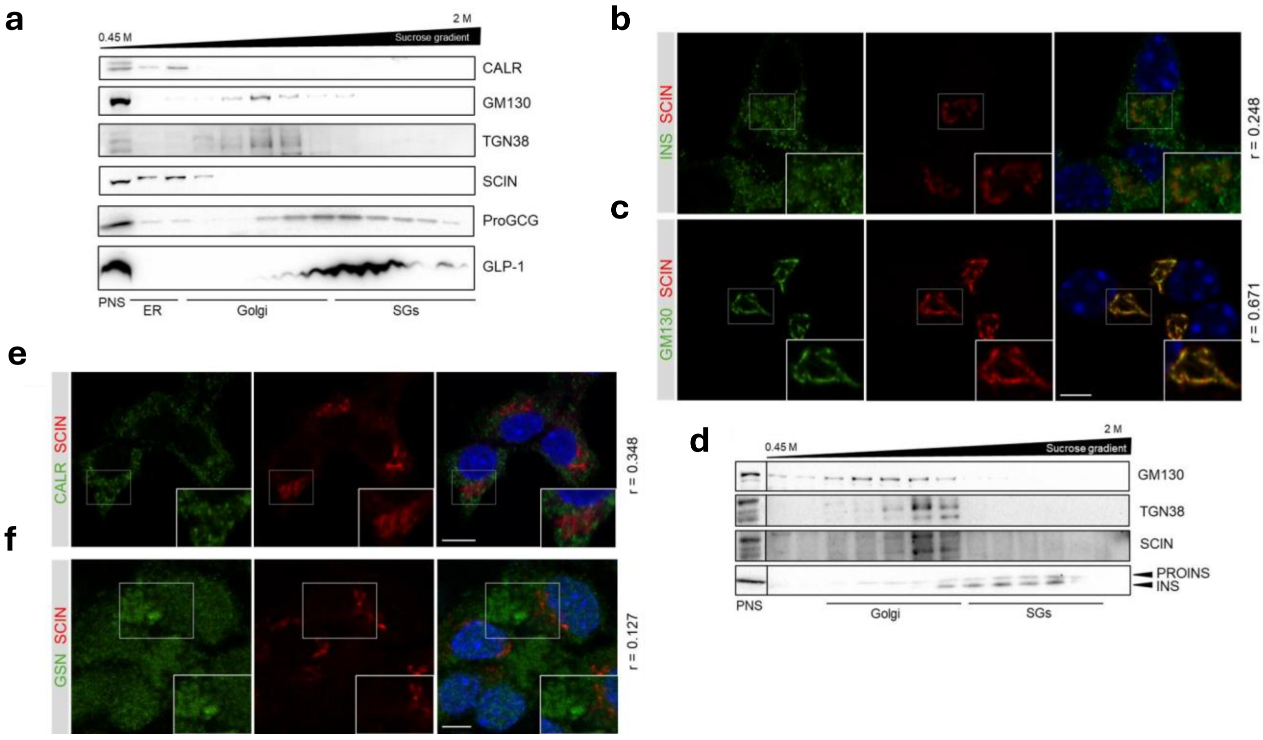


**Extended Data Figure 4. Golgi-restricted Scinderin distribution demonstrated by imaging and biochemical fractionation. a,** GLUTag cells were subjected to subcellular fractionation using a continuous sucrose gradient, fractions were analyzed by immunoblot. Calreticulin (CALR) is an endoplasmic reticulum (ER) marker, GM130 and TGN38 are Golgi markers. Proglucagon (ProGCG) is converted into GLP-1 and stored within secretory granules (SG). The post-nuclear supernatant (PNS) serves as a positive control, confirming the presence of proteins in the lysate prior to ultracentrifugation. Scinderin is present at the ER-Golgi interface. **b, c,** Representative immunofluorescence images of MIN6 cells co-stained for SCIN and other proteins. Insulin (b) shows partial colocalization with SCIN, whereas strong colocalization is observed with the Golgi marker GM130 (c). Average Pearson's correlation coefficients from colocalization analysis (n = 5) are shown. Scale bar, 5 μm. **d,** MIN6 cells were subjected to subcellular fractionation using a sucrose gradient, and fractions were analyzed by immunoblotting. GM130 and TGN38 serve as Golgi markers. Proinsulin (PROINS) is converted into insulin (INS) and stored in secretory granules (SG). The post-nuclear supernatant (PNS) serves as a positive control. Scinderin localizes to the Golgi body in mouse beta cells. **e,** Scinderin does not colocalize with ER protein Calreticulin (Calr). **f,** Scinderin and its closely related protein Gelsolin (GSN) show different localization patterns. Scale bar, 5 μm. Average Pearson's correlation coefficient from colocalization analysis (n = 5) is shown.

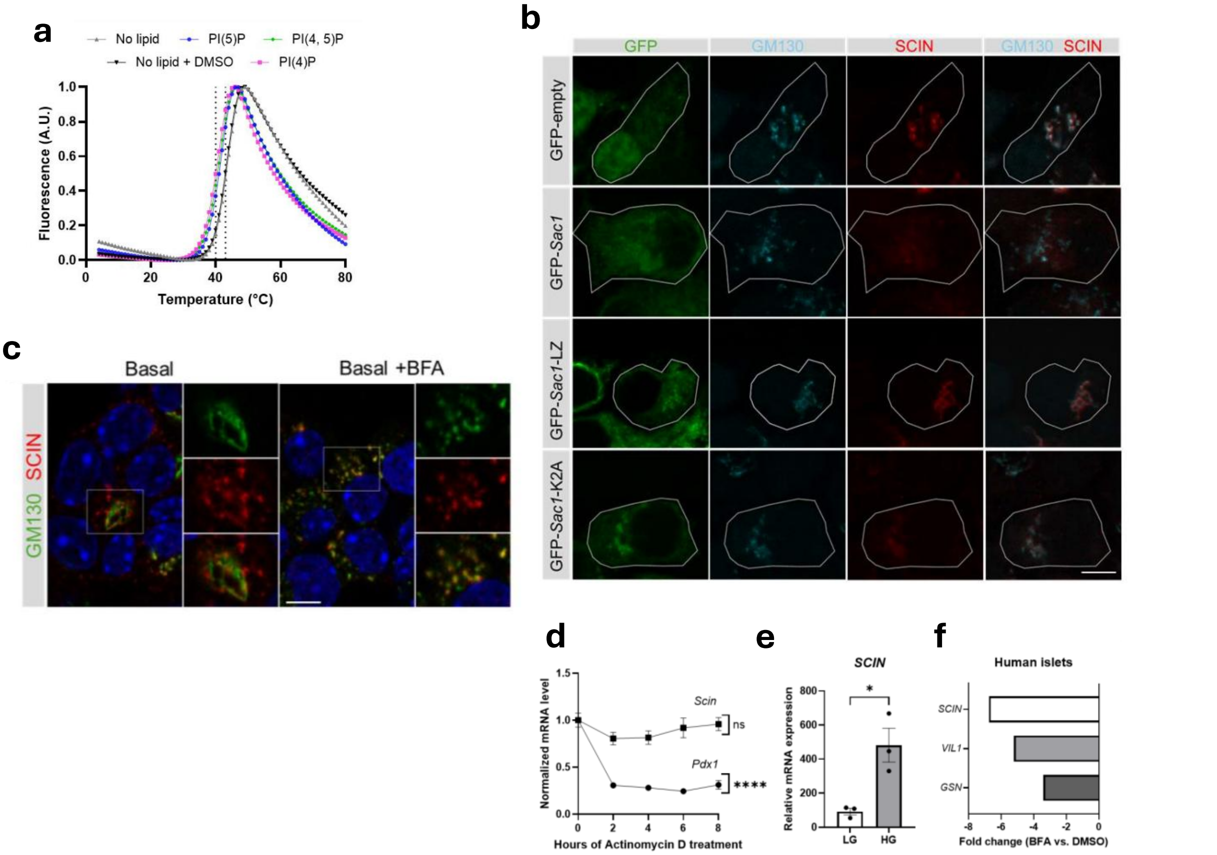


**Extended Data Figure 5. a,** Differential Scanning Fluorimetry confirmed binding to the top three PIPs, including Golgi-enriched lipid PI(4)P. **b,** GLUTag cells were transfected with different forms of phosphoinositide phosphatase SAC1, including wild-type (WT), LZ (ER-localized), and K2A (Golgi-localized) variants. Cells were stained for the Golgi marker GM130 and SCIN Wt-SAC1 and K2A-SAC1 depleted SCIN from the Golgi (quantification shown in Fig 2j). **c,** MIN6 cells were incubated under basal conditions with or without brefeldin A (BFA; 10 μM) for 1 hour to inhibit anterograde transport. SCIN appeared as vesicles budding from the Golgi apparatus during basal condition. Inhibition of Golgi trafficking caused SCIN vesicles to accumulate at the Golgi, demonstrating that these vesicles originate from this organelle. Scale bar, 5 μm. **d,** Scin mRNA stability was assessed in MIN6 cells following treatment with the transcriptional inhibitor actinomycin D. Pdx1 mRNA served as a positive control and decreased over time, whereas Scin mRNA remained relatively stable (n = 5). **e,** SCIN levels were analyzed by qPCR in EndoC-βH1 cells cultured under low glucose (LG, 5.6 mM) or high glucose (HG, 20 mM) conditions for 4 hours. Glucose increased SCIN expression (n = 3). Data are presented as mean ± SEM. *p ≤ 0.05, ****p ≤ 0.0001 vs. indicated groups by one-way ANOVA with Dunnett's post-hoc test (c), or unpaired t-test (d). **f,** Differentially expressed genes from human islets treated with brefeldin A were examined; among these several actin-remodeling proteins were identified. SCIN was downregulated in human islets under Golgi stress.

**a**

| Comparison | Up | Down | Total |
|---|---|---|---|
| KO1 vs. WT | 1 | 21 | 22 |
| KO2 vs. WT | 2 | 40 | 42 |
| Shared in KO | 0 | 16 | 16 |

**b**

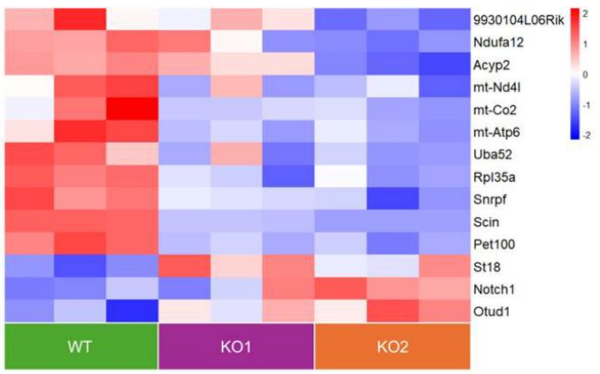


**c**

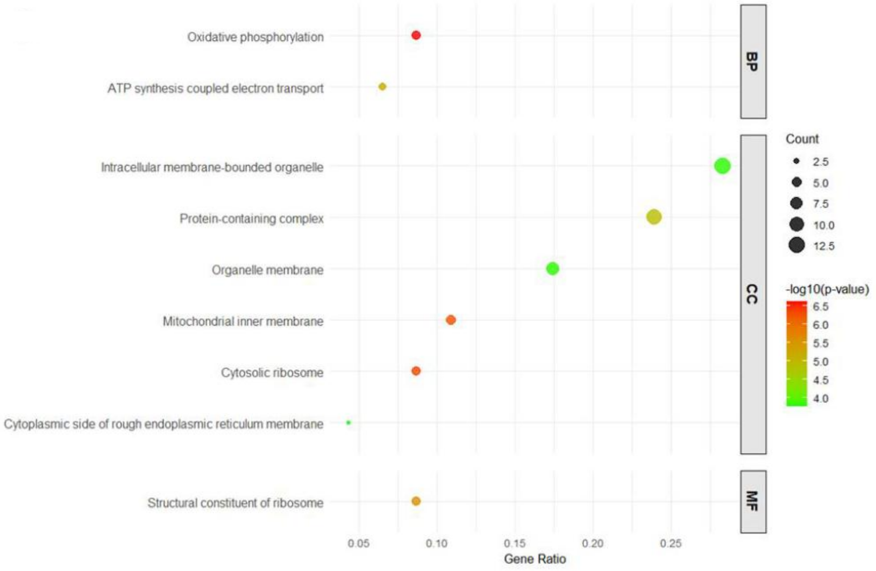


**Extended Data Figure 6. Transcriptomic changes upon Scinderin knockout in GLUTag cells. a,** Differentially expressed genes (DEGs) in knockout (KO) cells generated using different sgRNA, compared to control sgRNA (wild-type, WT). Shared DEGs among KOs are also shown. B) Heatmap of protein-coding differentially expressed genes in WT and KO cells. C) Enriched Gene Ontology (GO) terms in KO cells, categorized into biological process (BP), cellular component (CC), and molecular function (MF).

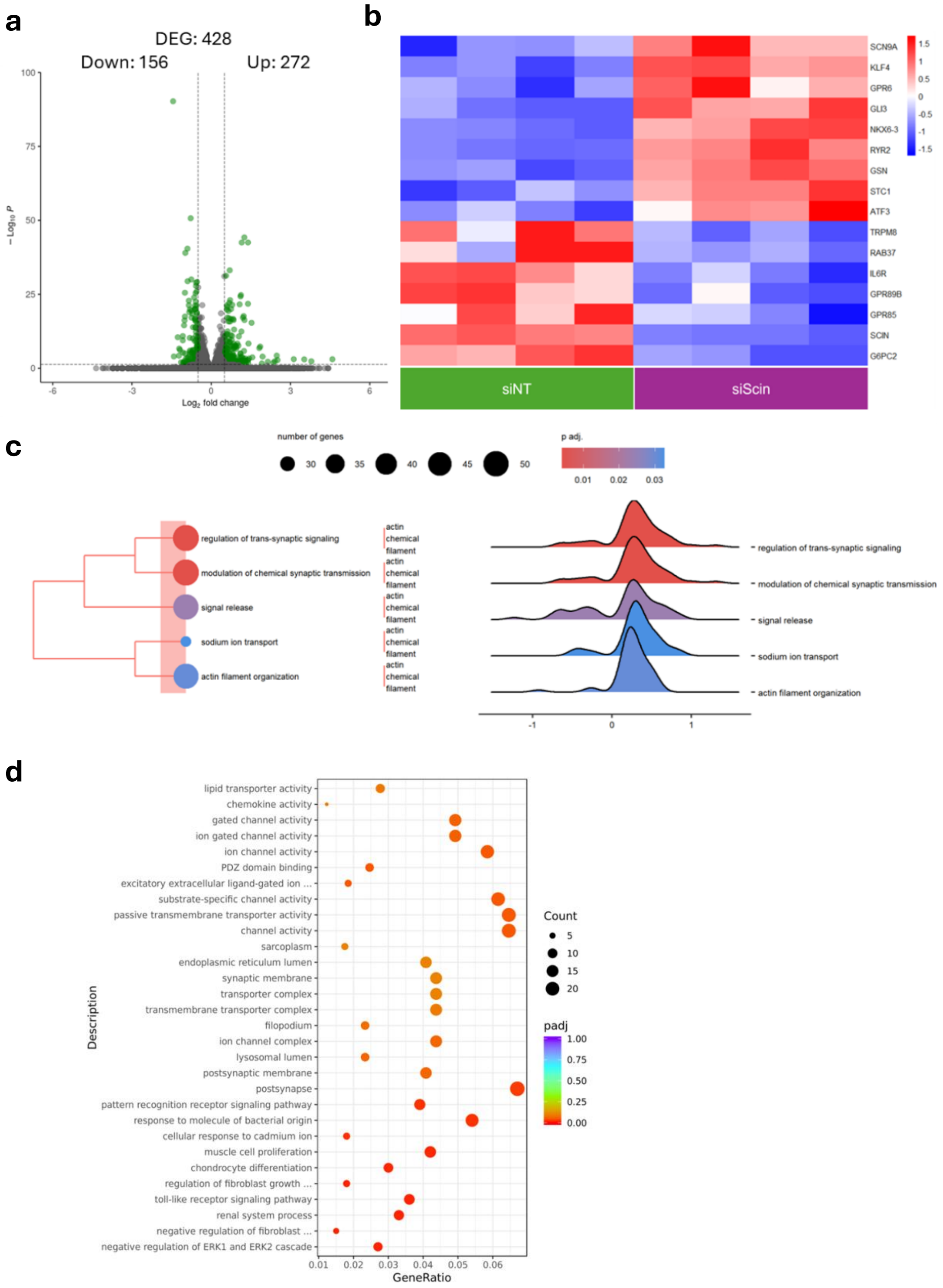


**Extended Data Figure 7. Transcriptomic alterations in EndoC-βH1 cells following Scinderin silencing.**
**a,** Volcano plot showing differentially expressed genes (DEGs) in EndoC-βH1 cells transfected with SCIN-targeting siRNA (siScin) compared to control siRNA (non-targeting, siNT). Green dots indicate significantly altered genes (p-adj ≤ 0.05 and |log2FC| ≥ 0.5), while grey dots represent non-significant changes in siScin vs siNT-transfected cells. **b,** Heatmap displaying selected protein-coding DEGs in siScin-silenced cells. **c,** Overrepresented biological processes identified through functional enrichment analysis of DEGs in siScin-silenced cells, shown as ridgeline plots where the X-axis represents the enrichment score and color indicates adjusted p-values. Overall, the analysis reveals widespread transcriptomic alterations in Scinderin-silenced EndoC-βH1 cells. **d,** Pathway enrichment analysis of deregulated genes in EndoC-βH1 cells following SCIN silencing. The dot plot illustrates significantly enriched Gene Ontology (GO) terms across the biological process, cellular component, and molecular function categories.

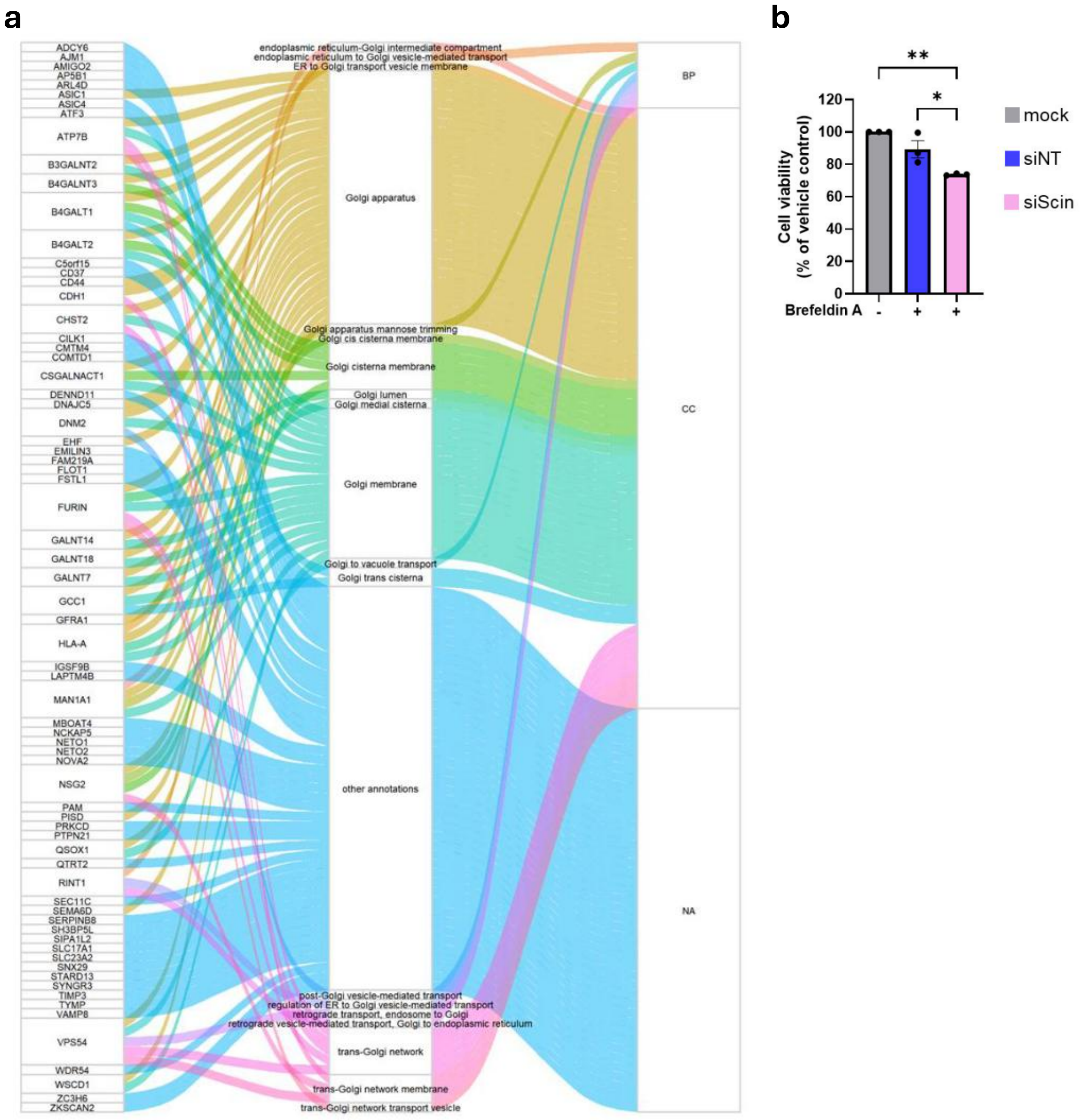


**Extended Data Figure 8. Scinderin-deleted human beta cells exhibit a Golgi-stress response. b,** Alluvial plot showing the 71 Golgi-associated DEGs from siScin-transfected EndoC-βH1 cells and their Gene Ontology categories (BP, biological process; CC, cellular component). **b,** EndoC-βH1 cells were treated with a low BFA dose (0.05 µg/mL), and cell viability was measured 16 hours later. SCIN-silenced EndoC-βH1 cells were more sensitive to the low BFA dose (n = 3) compared to siNT and untreated (mock) cells. Data are presented as mean ± SEM. *p ≤ 0.05, **p ≤ 0.01 vs. indicated groups by one-way ANOVA with Šidák's post-hoc test (C).

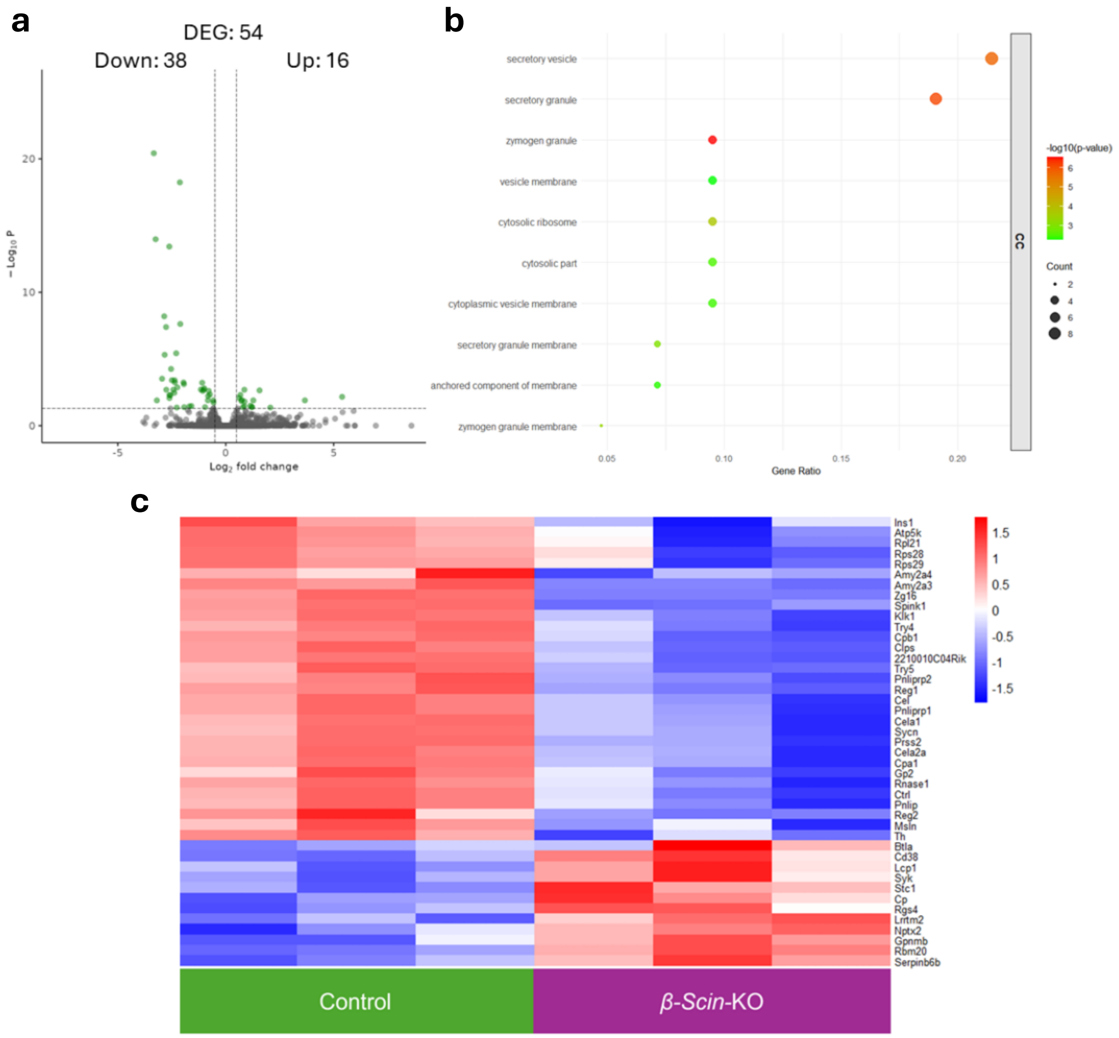


**Extended Data Figure 9. Transcriptomic changes in islets from β-Scin-KO mice. a,** Volcano plot showing differentially expressed genes (DEG) in β-Scin-KO islets. Green dots indicate significant changes in expression (p-adj ≤ 0.05 and |log2FC| ≥ 0.5); grey dots indicate non-significant changes in KO vs control islets. **b,** Dot plot of enriched Gene Ontology (GO) terms related to cellular compartment (CC) category in β-Scin-KO islets. **c,** Heatmap of protein-coding DEG in β-Scin-KO islets. Bulk RNA-sequencing results suggest a role for SCIN in vesicular trafficking.

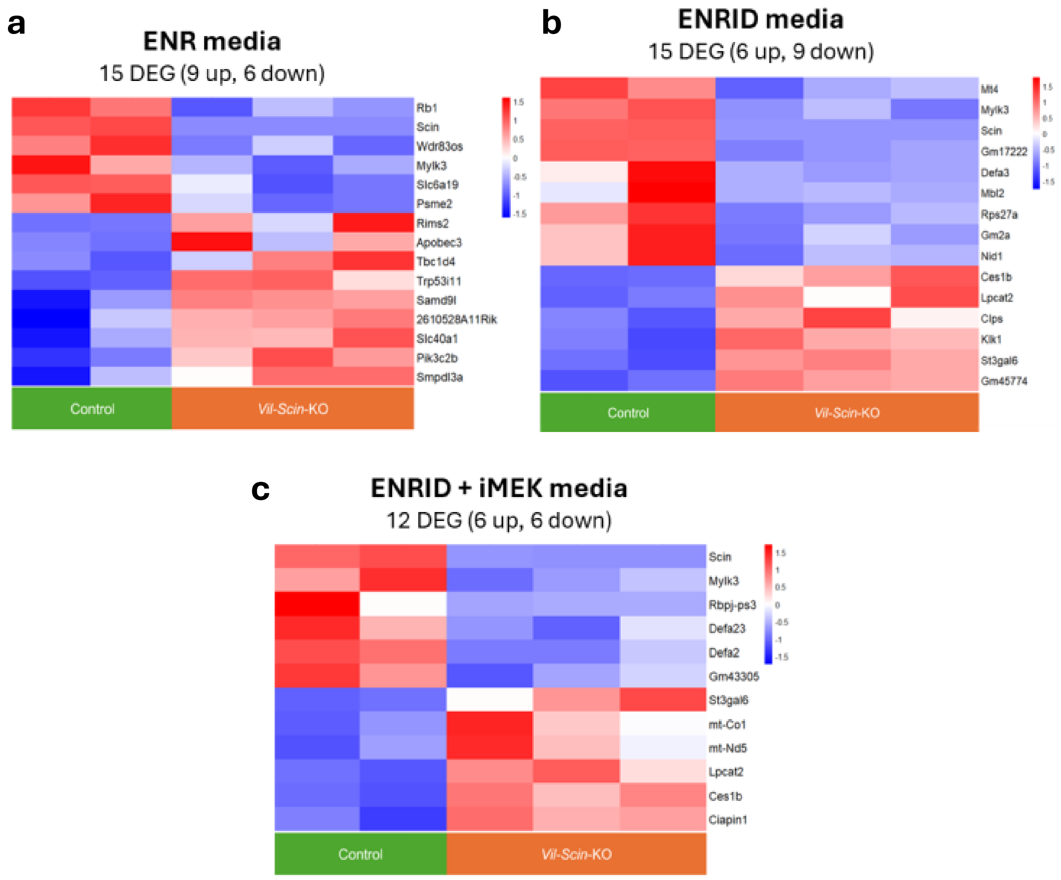


**Extended Data Figure 10. Transcriptomic alterations in intestinal Scinderin knockout mouse organoids. a-c,** Organoids were grown in distinct media conditions to enrich for specific cell types, followed by RNA isolation and RNA-seq. Heatmaps showing differentially expressed genes (DEGs) between control and Vil-Scin-KO organoids cultured in different media: **a,** ENR (standard culture media, all cell types), **b,** ENRID (goblet cells), and **c,** ENRID + iMEK (enteroendocrine cells).

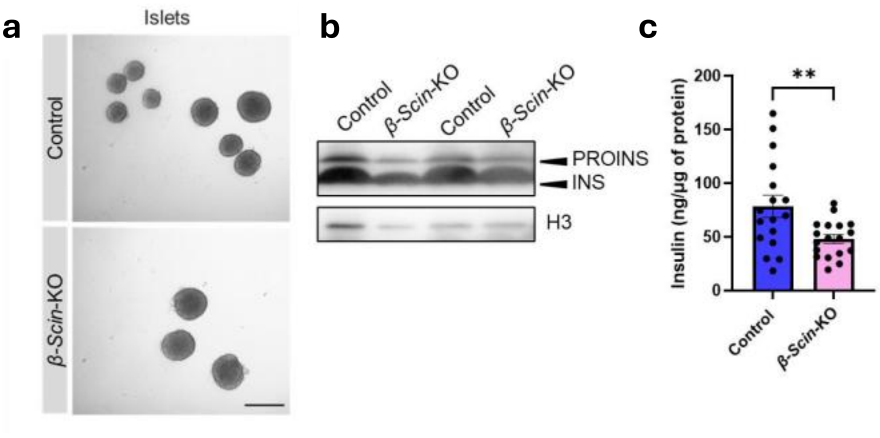


**Extended Data Figure 11. β-Scin-KO islets exhibit normal morphology but reduced insulin content.** **a,** Representative brightfield images of control and β-Scin-KO isolated islets showing comparable morphological features. Scale bar, 200 µm. **b,** (Pro)insulin content in the islets was assessed by immunoblotting. **c,** Insulin content was quantified by ELISA and normalized to total protein (n = 17–18). Both ELISA and immunoblot analyses revealed reduced (pro)insulin content in β-Scin-KO islets. Data are presented as mean ± SEM. **$p \leq 0.01$, by unpaired t-test

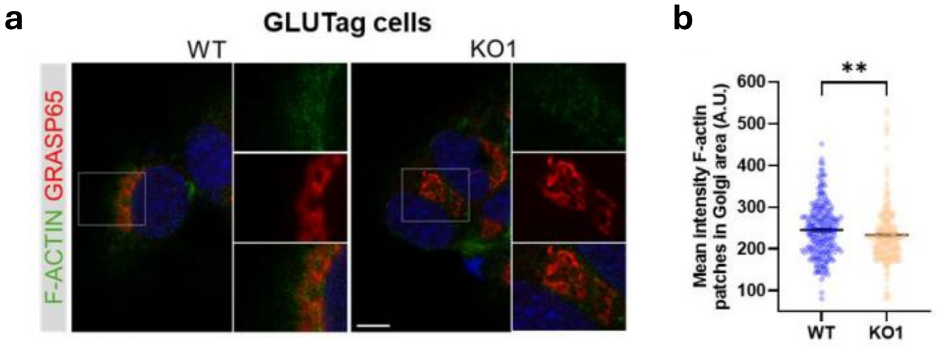


**Extended Data Figure 12. Altered actin dynamics in the absence of Scinderin**. **a,** Filamentous actin (F-actin) and the Golgi apparatus were immunostained in Scin knockout GLUTag cells. GRASP65 was used as a Golgi marker. **b,** The intensity of F-actin within the Golgi region was quantified for GLUTag (n = 219–231). Scale bar, 5 μm. The absence of Scin alters F-actin levels in the Golgi body. Data are presented as mean ± SEM. **p ≤ 0.01 vs. indicated groups by Mann-Whitney U test.

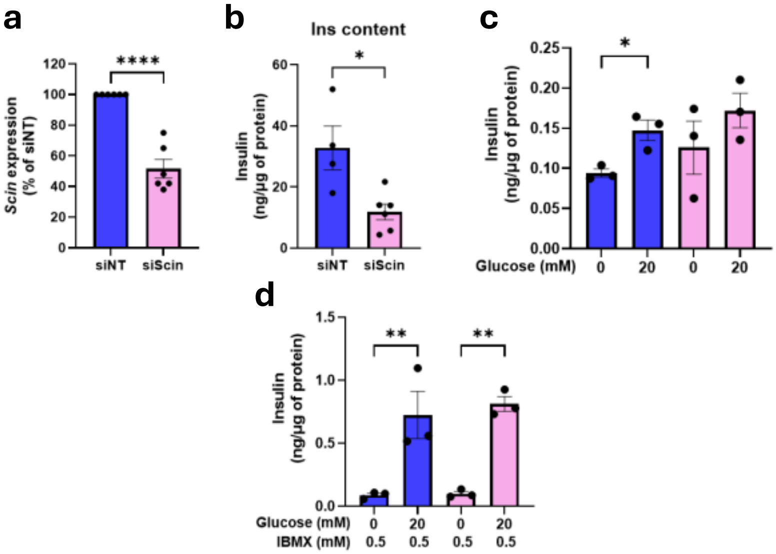


**Extended Data Figure 13. Silencing of Scin in MIN6 cells reduces insulin content and impairs glucose-stimulated secretion.** MIN6 cells were transfected with either non-targeting (siNT) or Scin-specific (siScin) siRNAs. **a,** Knockdown (KD) efficiency was validated by qPCR. **b,** Intracellular Insulin content was quantified by ELISA in siNT and siScin-silenced cells (n = 4–5). **c,** Insulin release was assessed under basal (0.5 mM glucose) and glucose-stimulated (20 mM) conditions (n = 3). Scin-KD cells failed to augment insulin secretion in response to glucose. **d,** Insulin secretion was measured under basal (0.5 mM glucose) and glucose-stimulated (20 mM) conditions in the presence of IBMX (0.5 mM) (n = 3). Scin KD cells demonstrated glucose responsiveness when IBMX was added. Data are presented as mean ± SEM. $*p \le 0.05$, $**p \le 0.01$, $****p \le 0.0001$ vs. indicated groups by unpaired t-test (a, b) or one-way ANOVA with Šidák's post-hoc test (c, d).

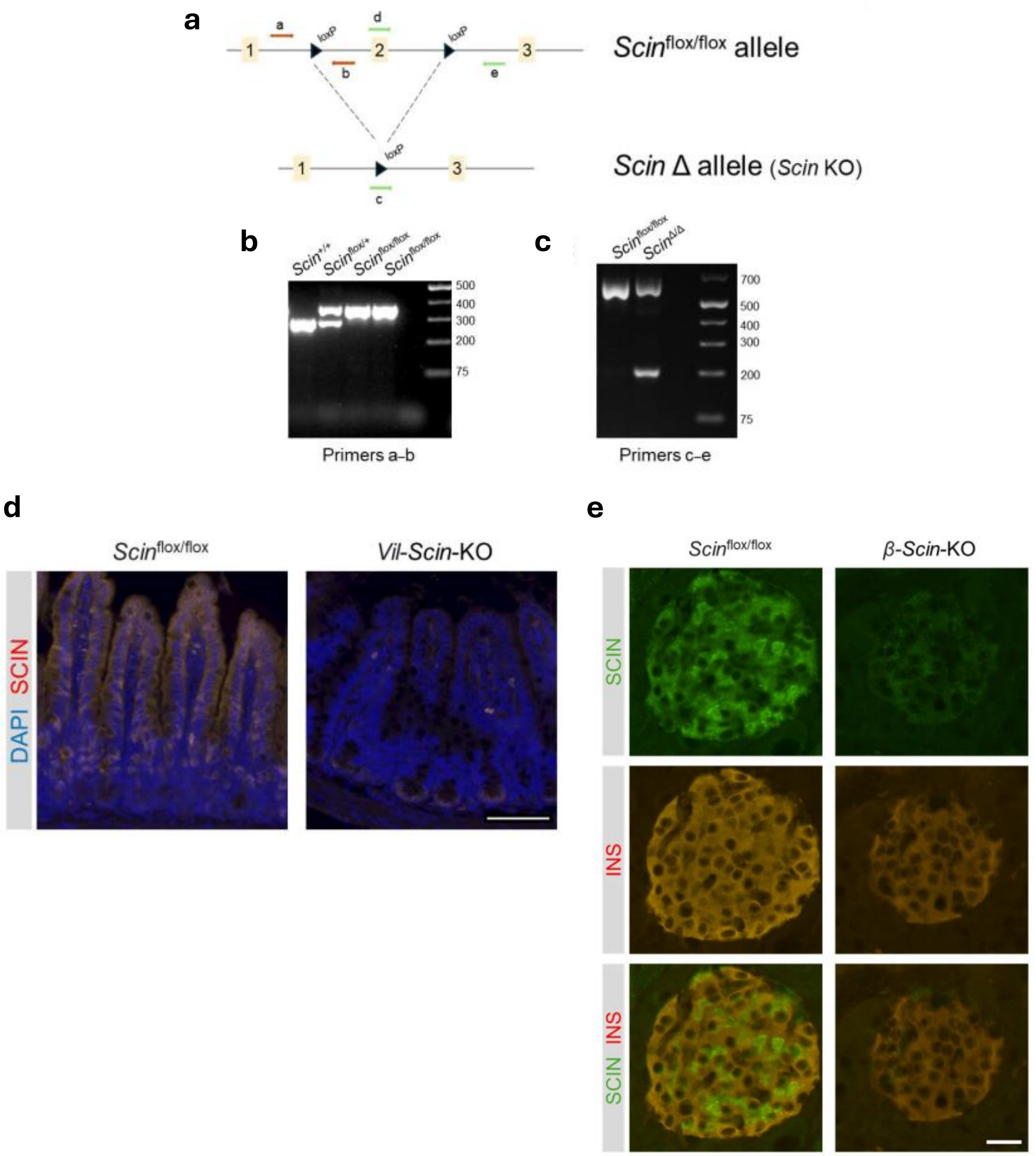


**Extended Data Figure 14. Scinderin KO mouse model. a,** Diagram showing Scinflox/flox and Scin deleted (Scin Δ) allele. Exons are indicated by yellow boxes. Exon 2 is conditionally targeted, loxP sites are shown. Upon crossing with tissue-specific Cre mice, exon 2 is excised in Cre-expressing tissues. Genotyping primers are indicated in red, while excision detection primers are shown in green. **b,** Mice are genotyped by PCR: wild-type (289 bp) and Scin flox (363 bp) allele. **c,** Scinderin excision is examined by PCR in Cre-expressing tissues. Deletion is confirmed by the appearance of a 210 bp band. **d,** Scinderin immunostaining on intestine from control (Scinflox/flox) and Vil-Cre x Scinflox/flox (Vil-Scin-KO) mice show loss of SCIN staining in the intestinal epithelium of KO mice. Scale bar, 100 μm. **e,** Scinderin beta cell KO mouse model. Immunostaining on pancreatic sections from control (Scinflox/flox) and Ins1-Cre x Scinflox/flox (β-Scin-KO) mice show loss of SCIN staining in beta cells from KO mice. Scale bar, 25 μm.

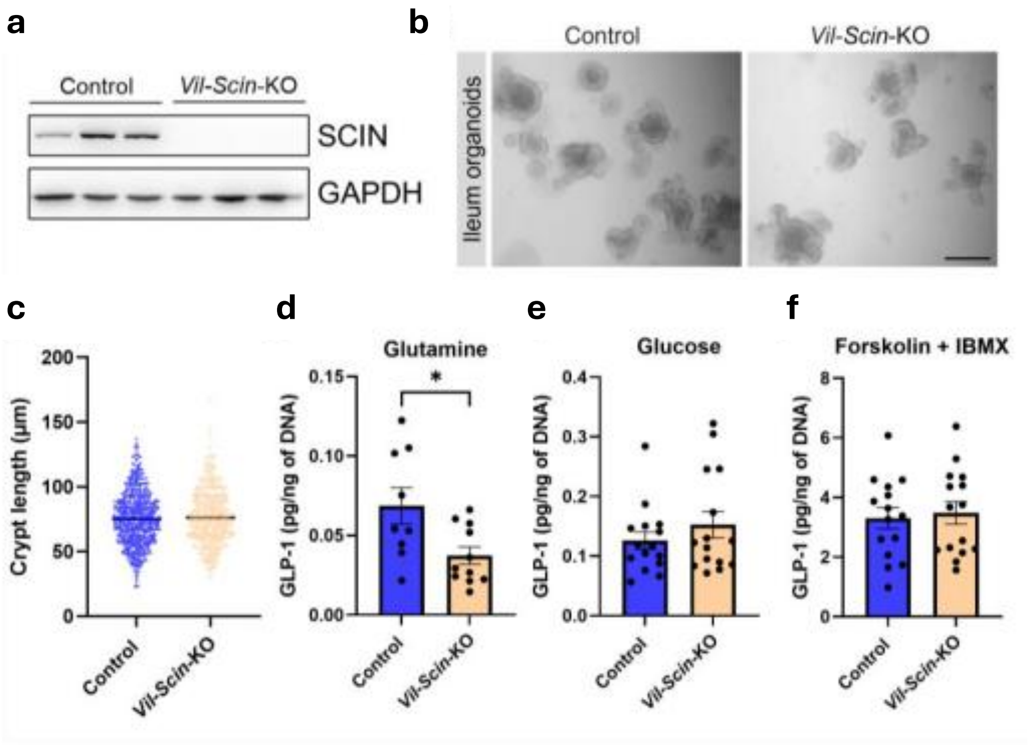


**Extended Data Figure 15. Organoids lacking SCIN exhibit normal morphology with a moderate defect in GLP-1 secretion. a,** Ileum-derived organoids were generated from control and Vil-Scin-KO mice. SCIN deletion was confirmed by immunoblot analysis of organoid protein lysates. **b,** Representative brightfield images of control and Vil-Scin-KO organoids cultured in ENR media showing comparable morphological features. Scale bar, 200 µm. **c,** Crypt length was measured in organoids, revealing no differences between control and Vil-Scin-KO (n = 3). **d-f,** GLP-1 secretion was evaluated following a 2-hour stimulation with different secretagogues: 2 mM glutamine (d), 17 mM glucose (e), and forskolin (10 µM) + IBMX (10 µM) (F). Vil-Scin-KO organoids displayed reduced GLP-1 secretion in response to glutamine, whereas no significant differences were observed upon stimulation with glucose or forskolin + IBMX (n = 3). Data are presented as mean ± SEM. *$p \leq 0.05$ by Mann-Whitney U test (c) or unpaired t-test (d–f).

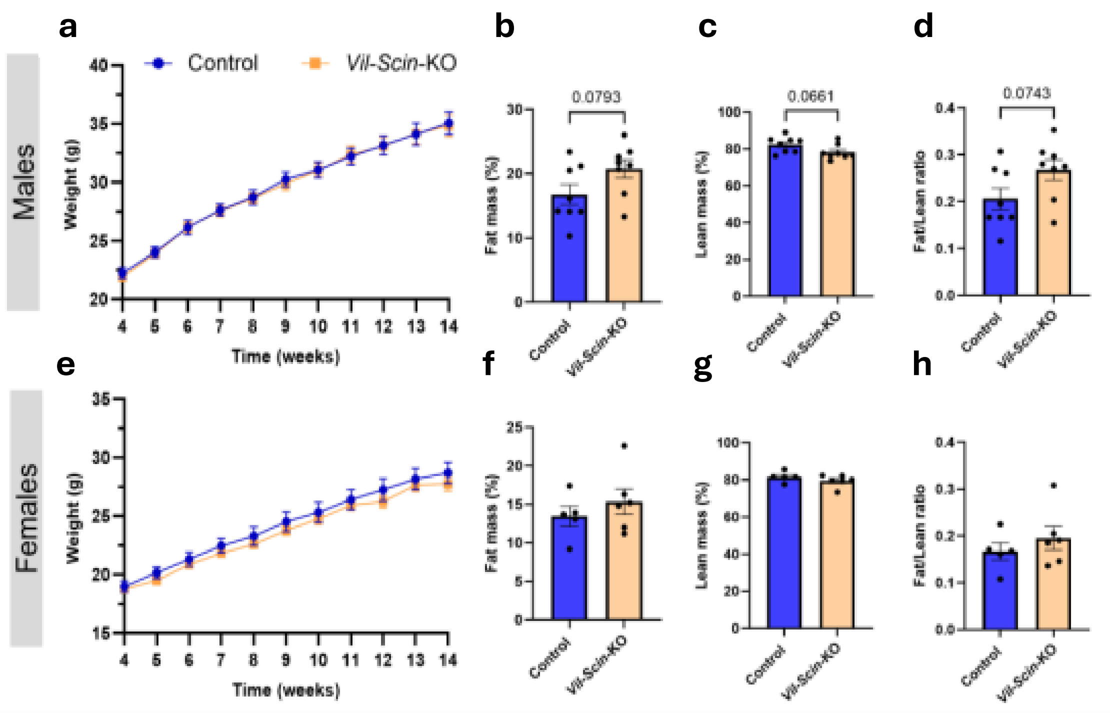


**Extended Data Figure 16. Longitudinal body weight and body composition of Vil-Scin-KO mice.** Body weight was measured weekly in male (**a**) and female (**e**) control and Vil-Scin-KO mice from 4 to 14 weeks of age (n = 10–15). No significant differences were observed between genotypes at any time point. Body composition was assessed at 5 months of age using magnetic resonance in males (n = 8) and females (n = 5–6). Fat mass (**b, f**), lean mass (**c, g**) and fat-to-lean mass ratio (**d, h**) are shown. Male Vil-Scin-KO mice displayed a tendency towards increased fat mass. Data are presented as mean ± SEM. Statistical analysis was performed using two-way ANOVA repeated-measures with Šidák's test (a, e) and by unpaired t-test (b–d, f–h).

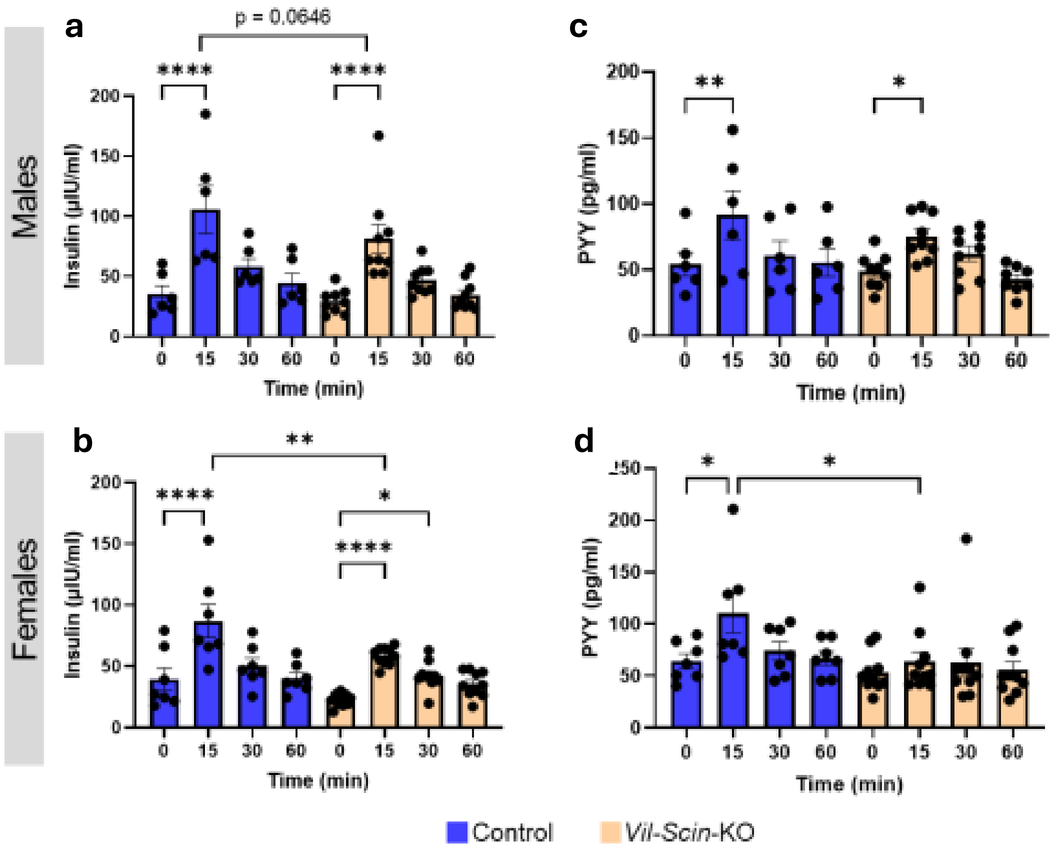


**Extended Data Figure 17. Insulin and PYY levels during oral glucose tolerance test in control and Vil-Scin-KO.** Male and female control and Vil-Scin-KO (16–19 weeks old) were fasted overnight before undergoing oGTT. **a, b,** Plasma insulin levels were measured from 0 to 60 minutes after glucose administration (n = 6–9). Both genders of Vil-Scin-KO showed a glucose-stimulated insulin response, though levels were modestly lower compared to those of control mice. **c, d,** PYY levels were also assessed over the same time period (n = 6–9). Male Vil-Scin-KO mice exhibited a blunted PYY secretion compared to controls, while female Vil-Scin-KO failed to increase PYY levels in response to glucose. Data are presented as mean ± SEM. *$p \le 0.05$, **$p \le 0.01$, ***$p \le 0.001$, ****$p \le 0.0001$ vs. indicated groups by one-way ANOVA repeated-measures with Šidák's post-hoc test.

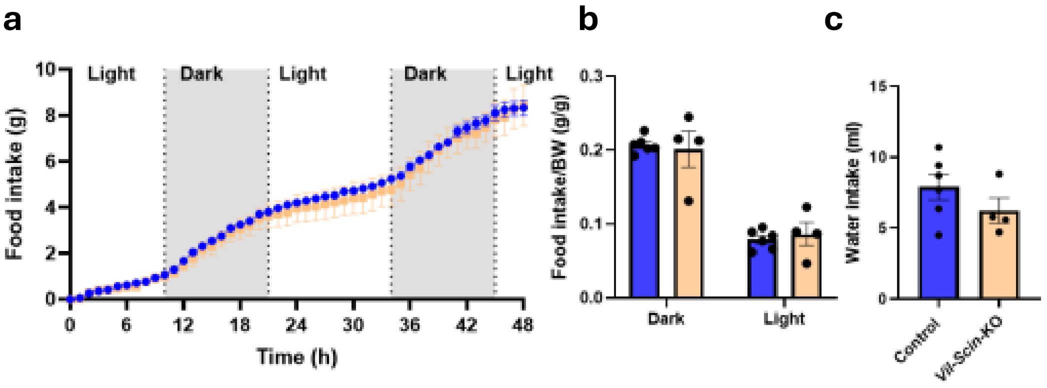


**Extended Data Figure 18. Food and water Intake of control and Vil-Scin-KO mice.**
**a,** Cumulative food intake over a 48-hour period (n = 4–6). **b,** Total food intake during dark and light phases, normalized to body weight. **c,** Total water intake over 48 hours (n = 4–6). Data are presented as mean ± SEM. *$p \leq 0.05$, vs. indicated groups by two-way ANOVA repeated-measures with Šidák's post-hoc test

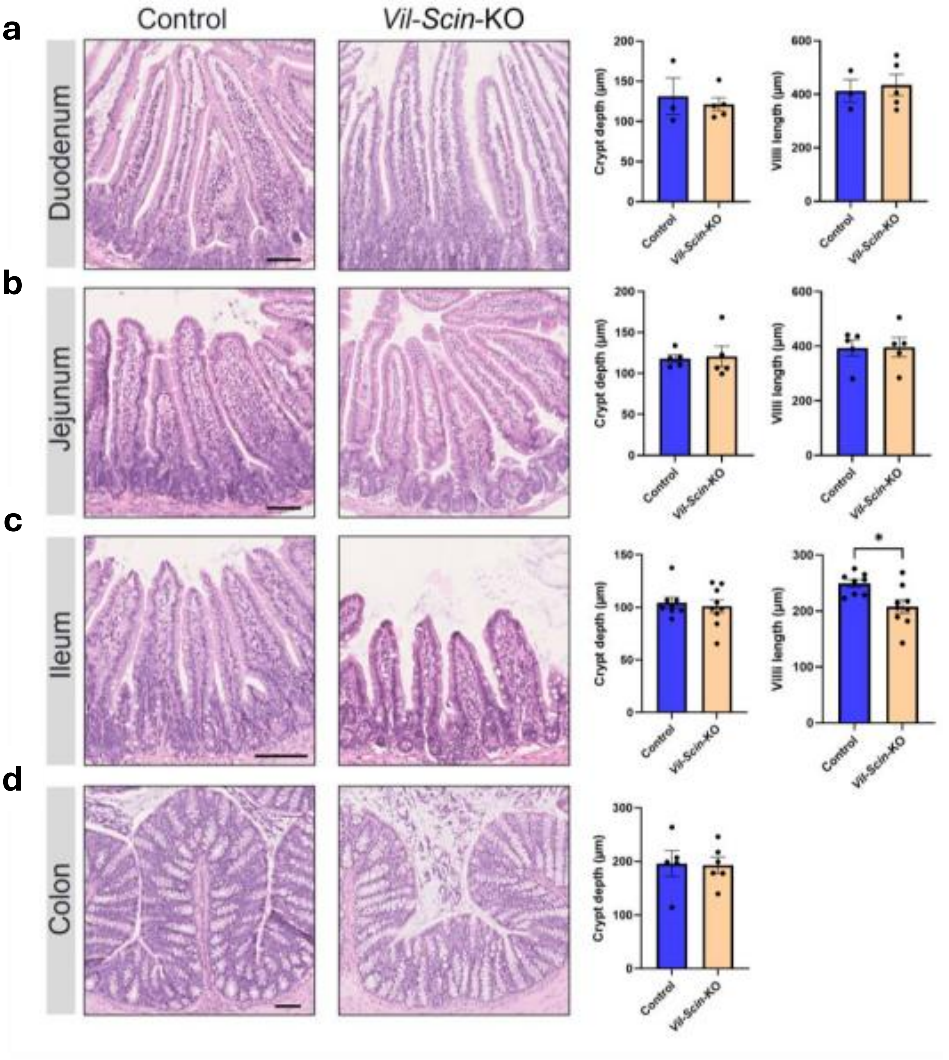


**Extended Data Figure 19. Histological examination of Vil-Scin-KO mice intestine.**

**a-d,** Representative H&E staining and corresponding morphometric analysis from control and Vil-Scin-KO intestine, including the duodenum (a), jejunum (b), ileum (c), and colon (d). Crypt depth and villi length were assessed across these intestinal regions (n = 3–9). No significant differences were observed between groups, except for a reduction in villi length in the ileum of Vil-Scin-KO mice. Scale bar, 100 µm. Data are presented as mean ± SEM. $^{*}p \leq 0.05$ by unpaired t-test.

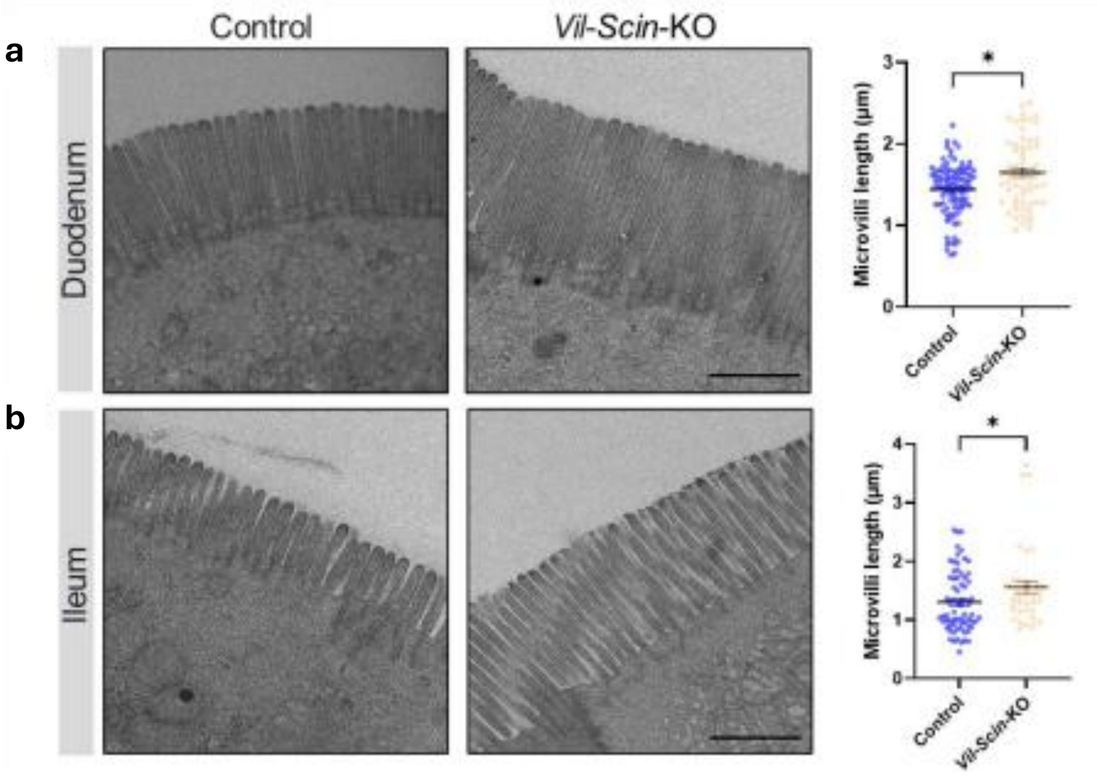


**Extended Data Figure 20. Electron micrographs of microvilli from control and intestinal Vil-Scin-KO mice. a, b,** Representative images and quantitative analyses of microvilli from the duodenum (a) and ileum (b) of control and Vil-Scin-KO intestine (n = 40–90 cells from 3–4 mice/genotype). Longer microvilli were observed in Vil-Scin-KO mice in both intestinal regions. Scale bar, 1 μm. Data are presented as mean ± SEM. *$p \le 0.05$ by Mann-Whitney U test.

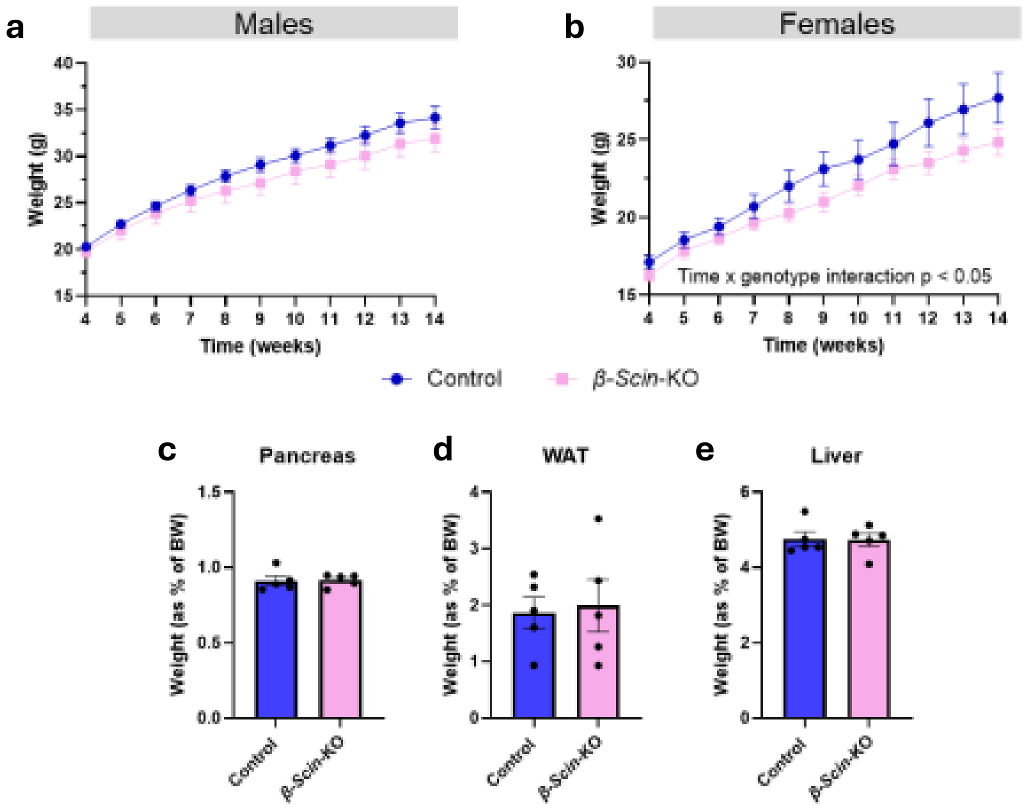


**Extended Data Figure 21. Longitudinal body weight and tissue weights in β-Scin-KO mice.** Body weight was measured weekly in male (**a**) and female (**e**) control and β-Scin-KO mice from 4 to 14 weeks of age (n = 8–13). No significant differences were observed between genotypes in males, whereas females showed genotype-dependent differences in weight gain over time. (**c–e**) At 20 weeks of age, female control and β-Scin-KO mice were sacrificed, and the weight of pancreas (**c**), visceral white adipose tissue (WAT) (**d**), and liver (**e**) were measured (n = 5). There were no differences between genotypes in the weight of these tissues. Data are presented as mean ± SEM. Statistical analysis was performed using two-way repeated-measures ANOVA with Šidák's test (a, b) and unpaired t-test or Mann-Whitney U test (c–e), as appropriate

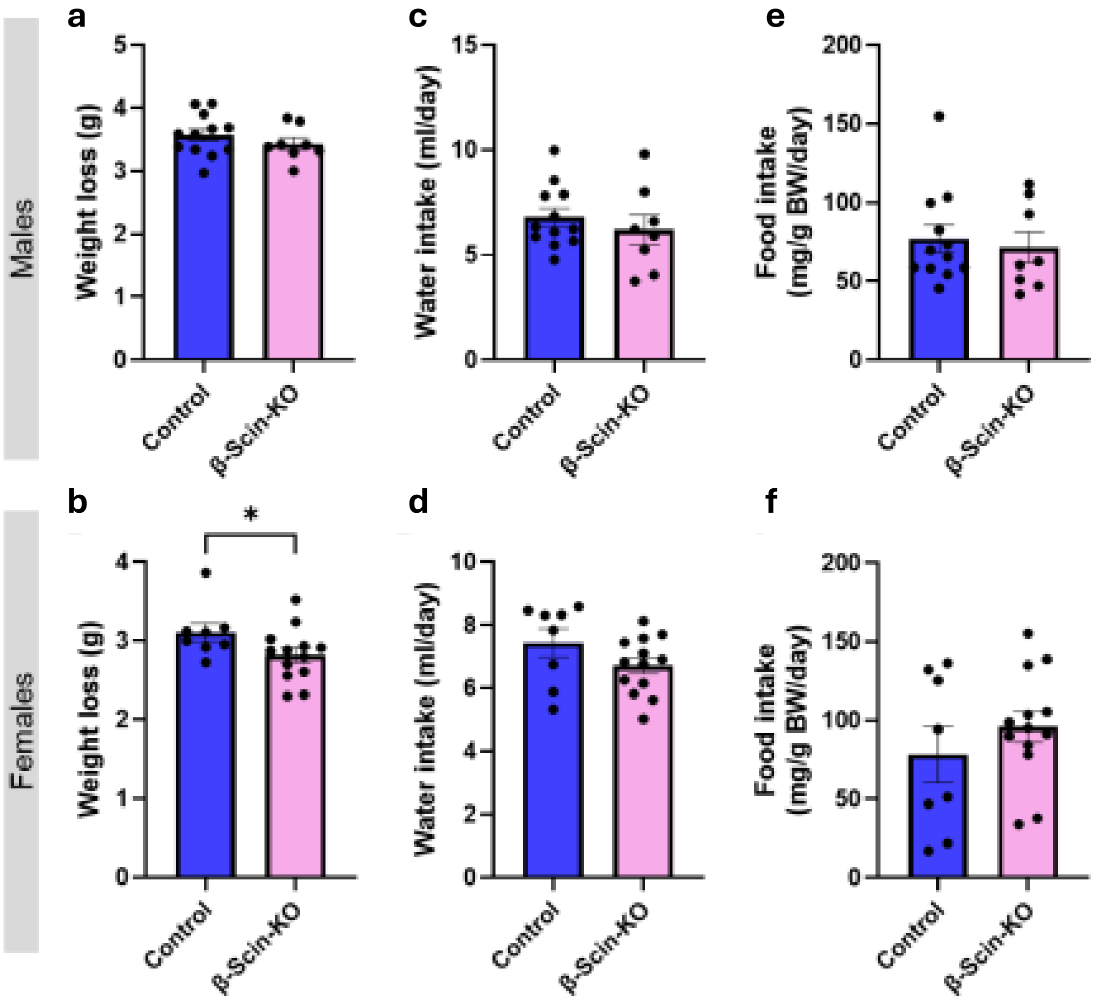


**Extended Data Figure 22. Weight loss and intake of control and β-Scin-KO mice during a fasting-refeeding experiment.** Food and water intake were measured in 15–17-week-old male and female mice after a 16-hour fast by monitoring chow and water weights at 4, 8, 24, and 48 hours following refeeding. **a, b,** Body weight loss after fasting in male (a) and female (b) control and β-Scin-KO (n = 8–13). No significant differences were observed in males, whereas KO females exhibited reduced weight loss. **c, d,** Daily water intake in male (c) and female (d) mice (n = 8–13). **e, f,** Daily food intake normalized to body weight in male (e) and female (f) mice (n = 8–13). No differences were observed in food or water intake. Data are presented as mean ± SEM. *$p \leq 0.05$, vs. indicated groups by unpaired t-test (A–E).

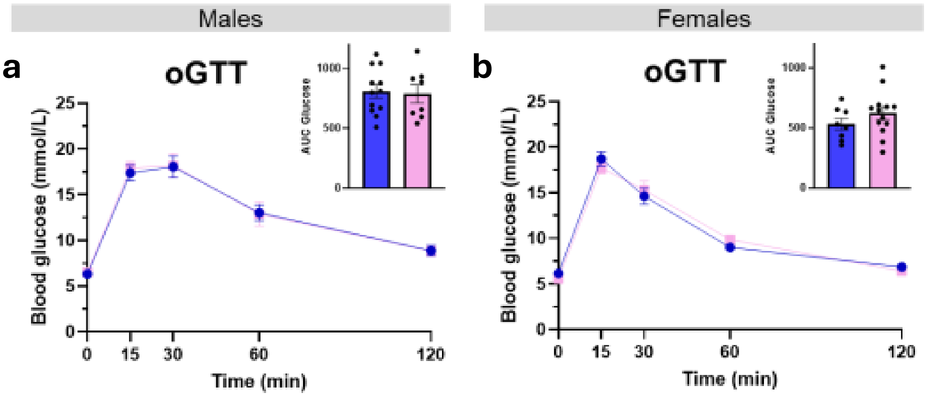


**Extended Data Figure 23. Glycemic responses during oral glucose tolerance tests (oGTT) in β-Scin-KO mice. a, b,** Male (a) and female (b) control and β-Scin-KO (16–19 weeks old) were fasted overnight prior to an oral glucose tolerance test (oGTT). Blood glucose levels and the corresponding area under the curve (AUC, inset) were measured over 0–120 min after oral glucose administration (n = 8–13).

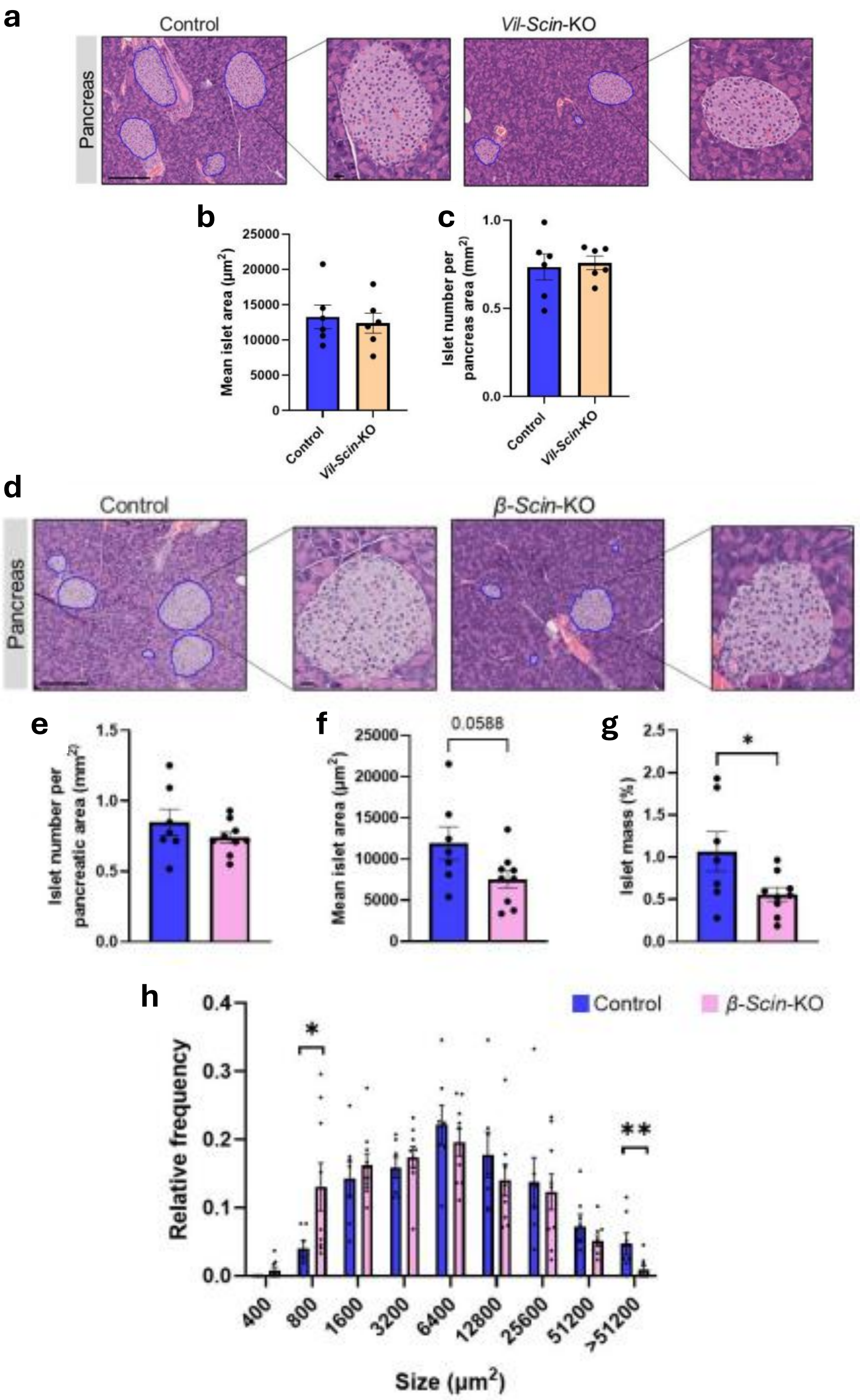


**Extended Data Figure 24. Histological analysis of pancreata from control, Vil-Scin-KO and β-Scin-KO. a,** Representative H&E- stained sections showing pancreatic islets from control and Vil-Scin-KO female mice. Scale bars: large bar, 200 μm; small bar, 20 μm. **b-c,** Morphometric analysis of islet size (b) and number (c). Data represent mean ± SEM from 6 mice per group (with >300 islets analyzed per group). No significant differences were observed between groups. Statistical analysis was performed using an unpaired t-test. **d,** Representative H&E-stained sections showing pancreatic islets from female control and β-Scin-KO mice. Scale bars: large bar, 200 μm; small bar, 20 μm. **e-g,** Morphometric quantification of islet number per pancreatic area (b), mean islet area (c), and total islet mass (d). No differences were observed in islet number, whereas β-Scin-KO exhibited reduced islet area and mass. **h,** Relative distribution of islet sizes. KO mice displayed a greater proportion of smaller islets and fewer large islets, consistent with their reduced islet mass. Data are presented as mean ± SEM from 6–9 mice per group (>350 islets analyzed per group). $^*p \leq 0.05$, $^{**}p \leq 0.01$ vs. indicated groups by unpaired t-test

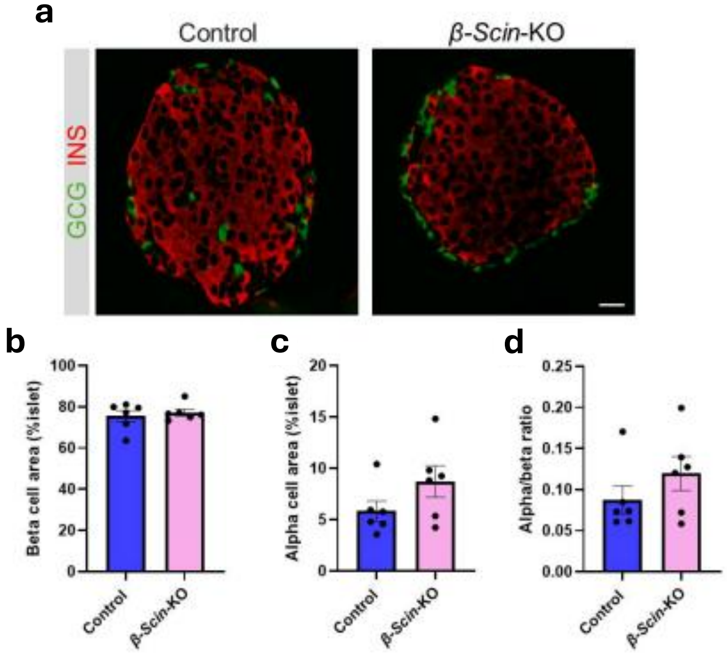


**Extended Data Figure 25. Analysis of beta and alpha cells in control and β-Scin-KO mice. a,** Representative immunofluorescence images of pancreatic islets stained for Glucagon (GCG, alpha cell marker) and Insulin (Ins, beta cell marker) in control and β-Scin-KO mice. Scale bar, 20 µm. **b-d,** Morphometric quantification of beta cell area (b), alpha cell area (c), and alpha-to-beta cell ratio (d). No differences were observed in beta cell area, alpha cell area or alpha-to-beta ratio between genotypes. Data are presented as mean ± SEM from 6 mice per group (80 islets analyzed per group). Statistical analysis was performed using unpaired t-test (b, c) and Mann-Whitney U test (D)

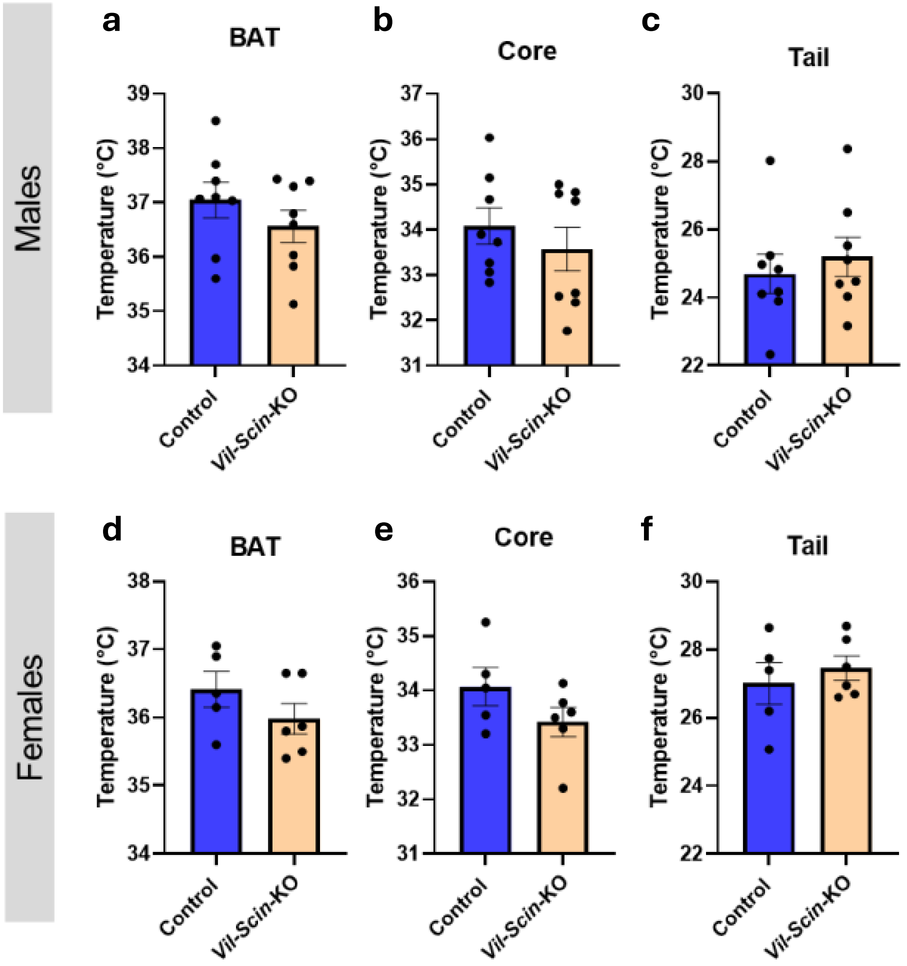


**Extended Data Figure 26. Body temperature measurements in control and Vil-Scin-KO mice.** Temperature was assessed at 5 months of age using an infrared camera in males (n = 8) and females (n = 5–6) mice. Temperatures of brown adipose tissue (BAT) (**a, d**), core (**b, e**), and tail (**c, f**) are shown. While differences did not reach statistical significance, KO mice of both sexes exhibited a similar directional trend across all tested regions. Data are presented as mean ± SEM. ns, not significant by unpaired t-test (a–b, d–h) and Mann-Whitney U test (c).

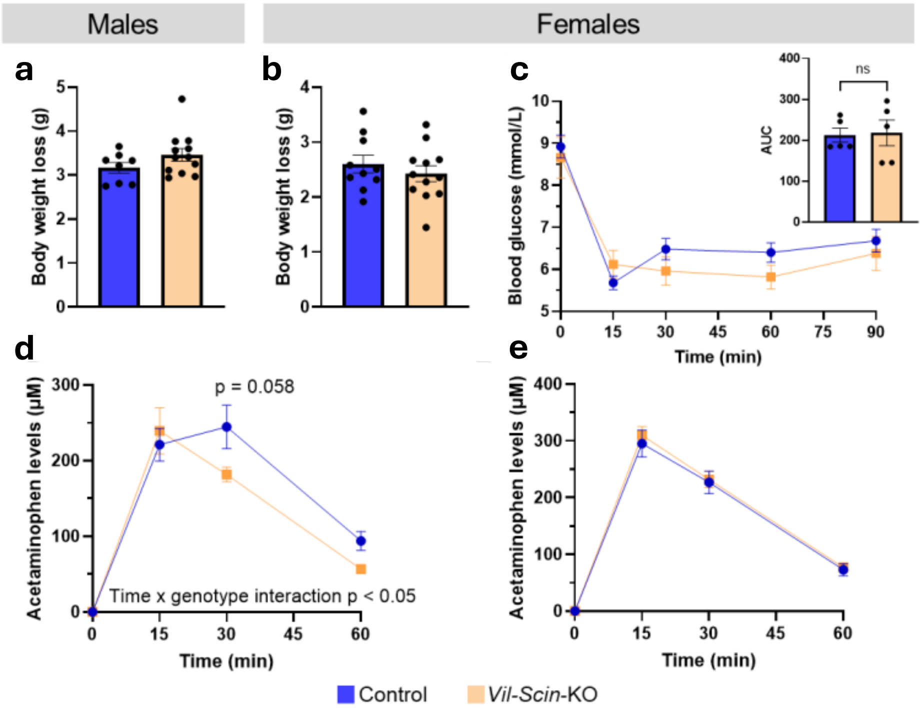


**Extended Data Figure 27. Weight loss after fasting, insulin tolerance test, and gastric emptying rate in control and Vil-Scin-KO mice. a, b,** Body weight loss after a 16-hour fast in male (a) and female (b) control and Vil-Scin-KO (16–19 weeks old; n = 8–12). No significant differences were observed between genotypes. **c,** Insulin tolerance test in 15-week-old female control and Vil-Scin-KO mice following a 2-hour fast. Blood glucose levels and area under the curve (AUC, inset) were measured over 0–90 min post-insulin injection (n = 5). Insulin sensitivity did not differ between groups. **d, e,** Gastric emptying assessed via levels of acetaminophen in plasma 0–60 min after oral co-administration of acetaminophen and glucose in males (d) and females (e) (n = 7–10). Acetaminophen levels decreased more rapidly in Vil-Scin-KO males (orange line), especially at 30 min. While no differences were observed in female mice. Data are presented as mean ± SEM. Unpaired t-test (A–B), Mann-Whitney U test (AUC of c), and two-way ANOVA repeated-measures with Šidák's test (c–e).

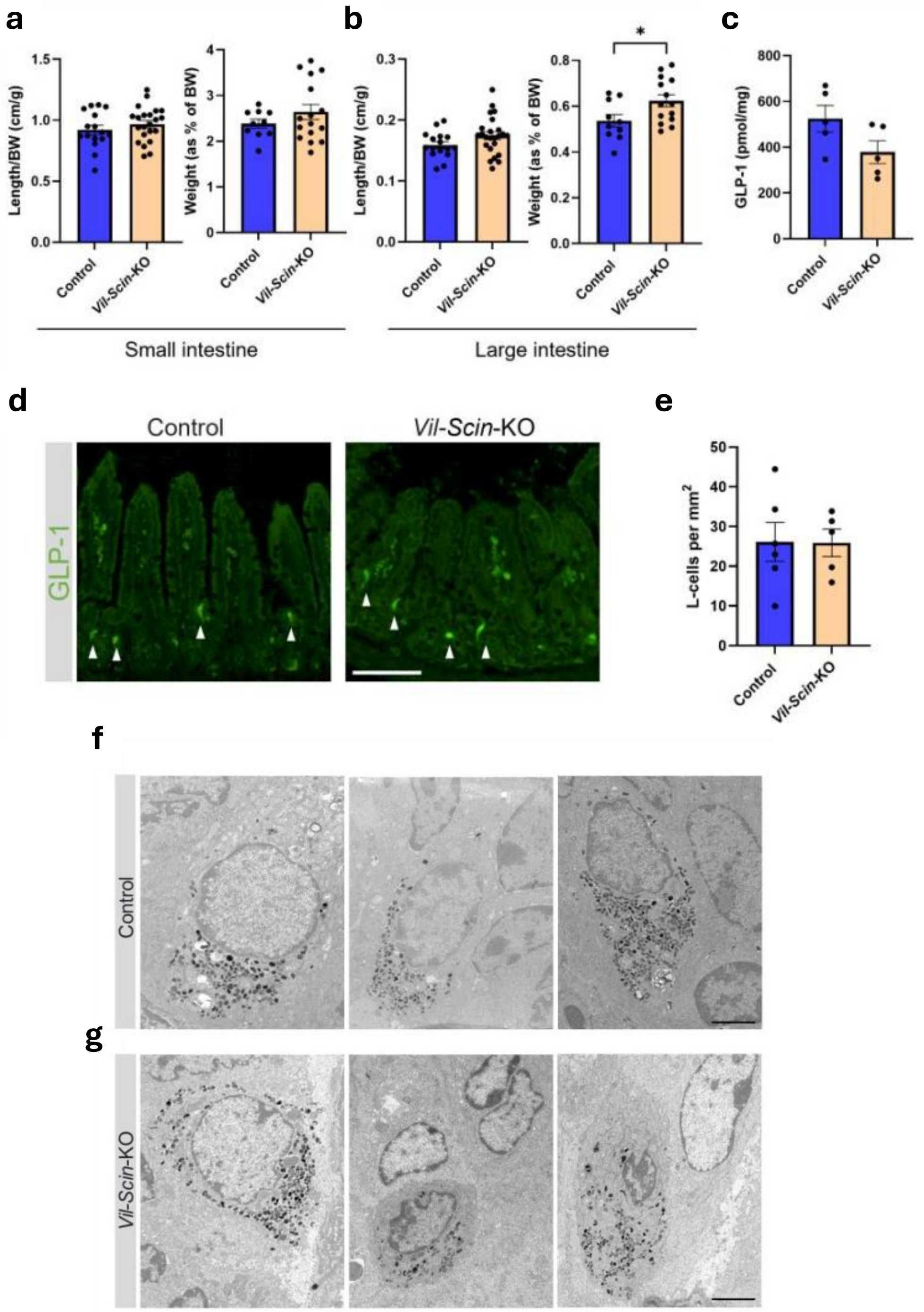


**Extended Data Figure 28. Macroscopic and microscopic characteristics, GLP-1 content, and number of L-cells in Vil-Scin-KO gut. a,** The small intestine was dissected from control and Vil-Scin-KO mice. Intestinal length (left) and weight (right) relative to body weight (BW) were measured (n = 10–22). **b,** Same analysis as in (a), but for the large intestine (n = 10–24). An increased large intestine weight was observed in Vil-Scin-KO mice. **c,** GLP-1 content was quantified in the ileum and normalized per mg of tissue (n = 5). **d,** Representative immunofluorescence image of ileum stained for GLP-1. Arrowheads indicate L-cells. Scale bar, 100 µm. **e,** Quantification of GLP-1 positive cells, presented as the number of L-cells per $mm^2$ of intestine epithelium (n = 5–6). Data are presented as mean ± SEM. ns, not significant, *p ≤ 0.05 by unpaired t-test. **f, g,** Electron micrographs of enteroendocrine cells of intestinal Scinderin KO mice. Representative images of ileal enteroendocrine cells from control (f) and Vil-Scin-KO intestine (g). Scale bar, 2 µm.

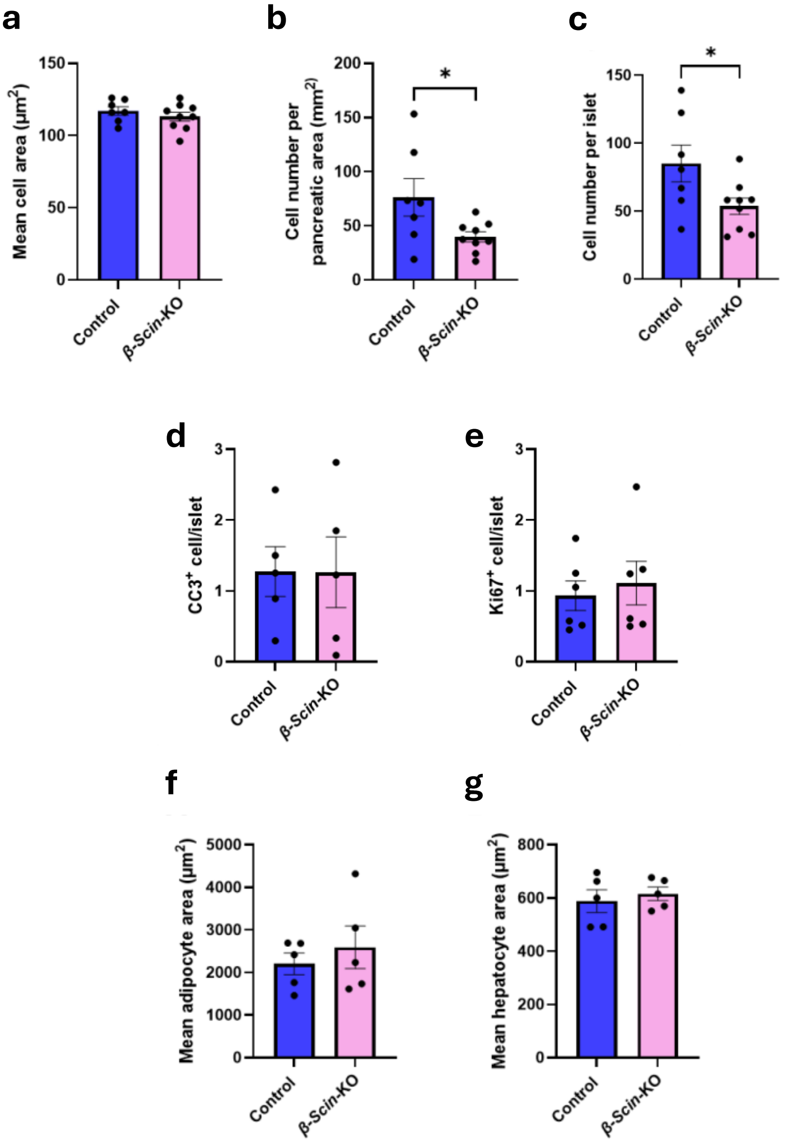


**Extended Data Figure 29. Histological and cell turnover analysis of pancreatic islet cells and histometry of adipocytes and hepatocytes from control and β-Scin-KO.** Morphometric analysis of islet cell parameters: mean area of individual islet cells (**a**), number of islet cells per pancreatic area (**b**), and number of islet cells per islet (**c**). No differences were observed in individual cell area, whereas β-Scin-KO islets showed a reduced number of cells per pancreatic area and per islet. Data are presented as mean ± SEM from 6–9 mice per group (>350 islets analyzed per group). *$p \leq 0.05$ vs. indicated groups by unpaired t-test. **d, e,** Analysis of cell turnover in islets from β-Scin-KO mice. Pancreatic sections were co-stained with insulin and cleaved-caspase 3 (CC3, a) or Ki67 (b) to assess apoptosis and proliferation, respectively, in control and β-Scin-KO islets. No differences were observed between genotypes. Data are presented as mean ± SEM from 5 mice per group (>155 islets analyzed per group). Statistical analysis was performed using unpaired t-test. igure S15. **f, g,** Adipocyte and hepatocyte size in β-Scin-KO mice. White adipose tissue (WAT) and liver were collected for histological analysis, and cell areas were quantified from tissue sections. Mean adipocyte area (f) in visceral WAT of control and in β-Scin-KO female mice (n = 5) and mean hepatocyte area (g) in the liver of control and in β-Scin-KO female mice (n = 5). No differences were observed in the adipocyte or hepatocyte size between groups. Data are presented as mean ± SEM. Statistical analysis was performed using unpaired t-test (A, B).

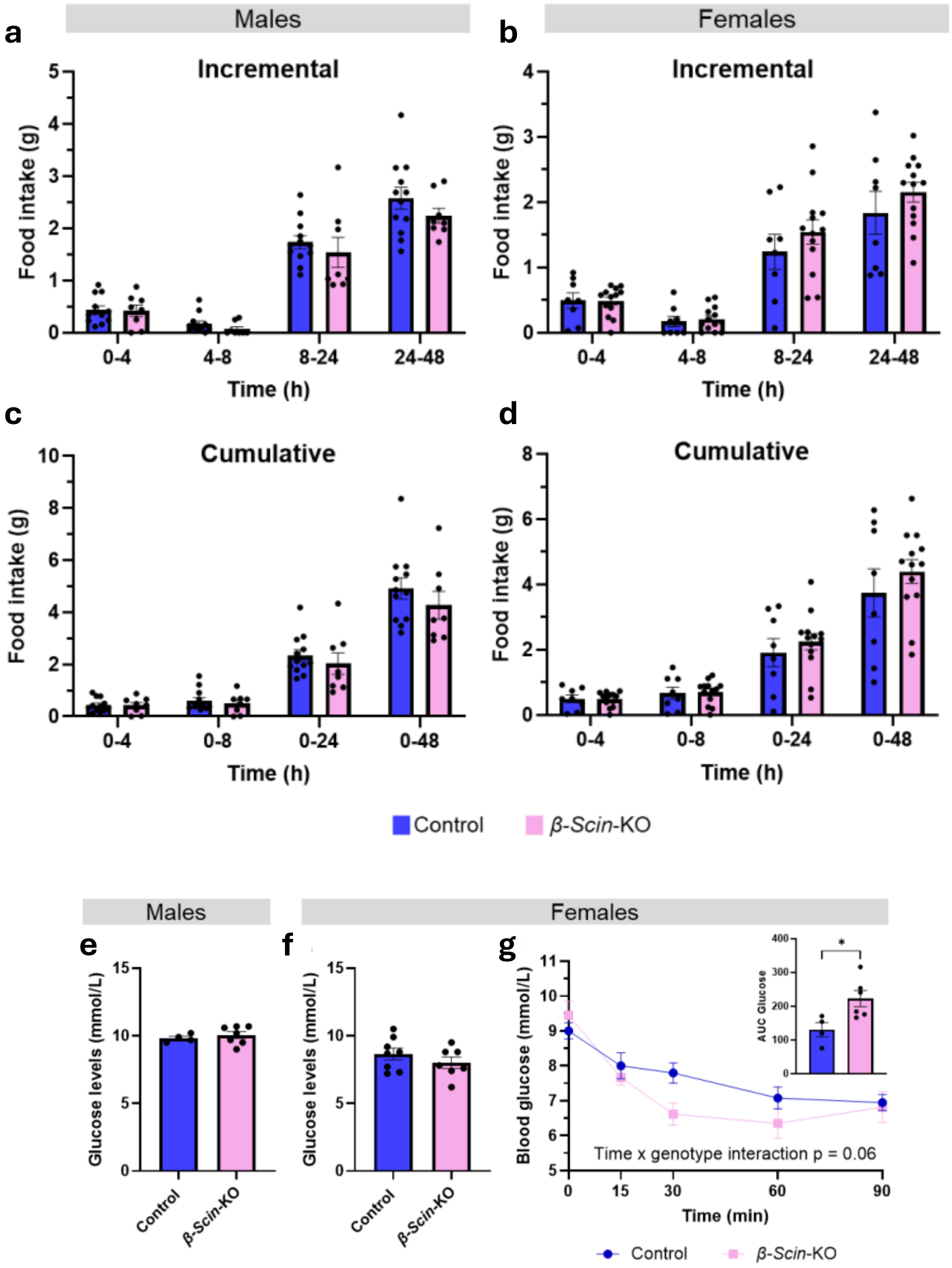


**Extended Data Figure 30. Incremental and cumulative food intake during a fasting-refeeding and glucose homeostasis and insulin tolerance of control and β-Scin-KO mice experiment.** Male and female mice (15–17-weeks old) were fasted overnight (16 h), after which a pre-weighed amount of chow was reintroduced. Food intake was assessed by reweighing the chow at 4, 8, 24, and 48 hours. **a, b,** Incremental food intake in males (a) and females (b) during the experiment (n = 8–13). **c, d,** Cumulative food intake in males (c) and females (d). No differences in food consumption were observed between genotypes in either sex. Data are presented as mean ± SEM. Statistical analysis was performed using two-way repeated-measures ANOVA with Šidák's test (a–d). **e, f,** Glucose homeostasis and insulin tolerance in β-Scin-KO mice. Random-fed blood glucose levels of male (a) and female (b) control and β-Scin-KO mice (12–16 weeks old; n = 4–8). No differences were observed between genotypes. **c,** Female mice (14 weeks old) were fasted for 2 hours and subjected to an insulin tolerance test (ITT). Blood glucose levels and the corresponding area under the curve (AUC, inset) were measured over 90 min following insulin administration (n = 4–6). β-Scin-KO mice exhibited greater reduction in blood glucose during ITT, indicating increased insulin sensitivity. Data are presented as mean ± SEM. *$p \leq 0.05$ vs. indicated groups by two-way repeated-measures ANOVA with Šidák's test (g) and unpaired t test (e, f, AUC of g)

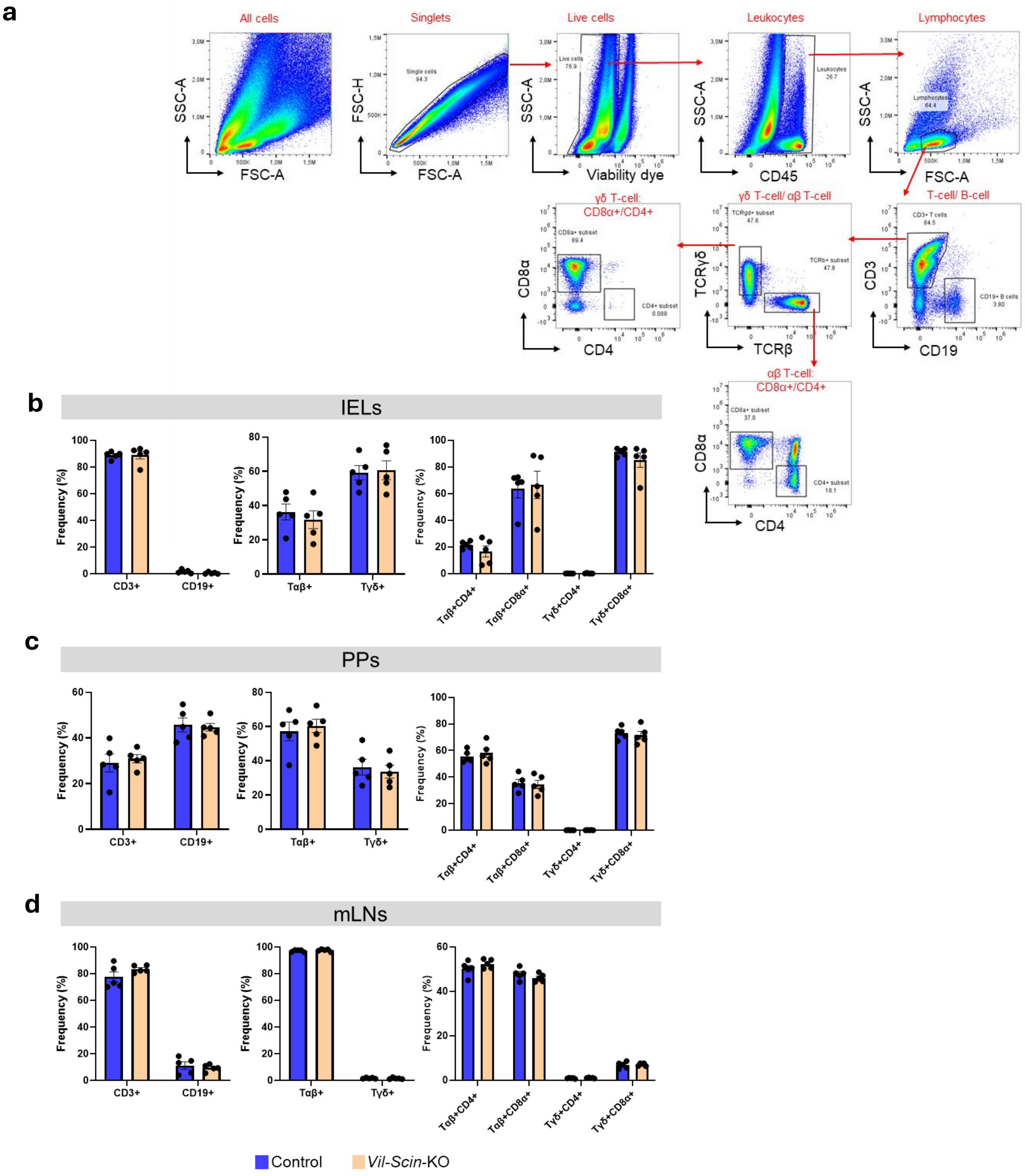


**Extended Data Figure 31. Flow cytometry gating strategy and analysis of lymphocyte populations in intestinal tissues in control and Vil-Scin-KO mice. a,** Single-cell suspensions from mesenteric lymph nodes, Peyer's patches, and intraepithelial lymphocytes were stained with a viability dye and fluorescently labeled antibodies, then analyzed by flow cytometry. This approach enabled the identification of T and B cells, as well as subsets of αβ and γδ T cells, including their respective CD4+ and CD8α+ populations. Single-cell suspensions were prepared from intraepithelial lymphocytes (IELs) (**b**), Peyer's patches (PPs) (**c**), and mesenteric lymph nodes (mLNs) (**d**). Panels show the frequency of lymphocyte subsets relative to total lymphocytes (left), CD3+ cells (middle), and Tαβ and Tγδ subpopulations as percentage of total Tαβ or Tγδ cells (right). No significant differences were observed between genotypes in any of the tissues. Data represent mean ± SEM from 5 mice per group. Statistical analysis was performed using an unpaired t-test or Mann-Whitney U test, as appropriate.

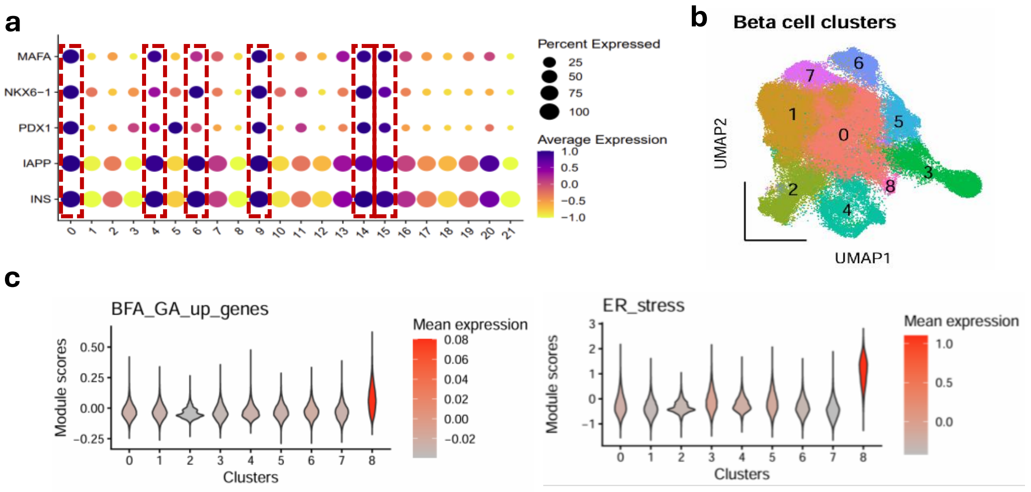


**Extended Data Figure 32. Identification of a BFA-responsive and ER-stress–associated β-cell state in healthy, prediabetic, and T2D islets. a,** To investigate the role of Scin in T2D, we analyzed publicly available scRNA-seq data (GSE221156) from healthy, prediabetic and T2D pancreatic islet cells. Cells were clustered using Seurat, and β-cells were identified based on canonical markers (MAFA, NKX6-1, PDX1, IAPP, INS). **b,** nine distinct β-cell subclusters were identified and **c,** GA genes upregulated upon BFA treatment in islets (padj < 0.01 and log2FC > 4), along with canonical ER stress markers (DIT3, HSPA5, ATF4, and XBP1), were enriched in cluster 8.
