## Supplemental Methods for "Scinderin-driven Golgi Actin Remodeling coordinates GLP-1 and insulin secretion to regulate glucose homeostasis"

Mice

All animal procedures were approved by the Finnish National Animal Experiment Board (ESAVI/5824/2018, ESAVI/21996/2021, ESAVI/33102/2024) and conducted in the preclinical animal facility at Tampere University under standard 12‑h light–dark cycles with *ad libitum* access to chow and water. Scin^flox/flox^ mice, generated by flanking exon 2 of *Scin* with loxP sites, were gifted from Michael Glogauer. *Vil1‑Cre* (JAX 021504) and *Ins1‑Cre* (JAX 026801) mice were used to generate intestinal (*Vil‑Scin‑KO*) and β‑cell (*β*‑*Scin‑KO*) knockouts, respectively. Cre and floxed alleles were genotyped using PCR (Extended Data Table 1). Animals were maintained on a C57BL/6J background and age‑ and sex‑matched littermates were used as controls. No differences were observed between floxed and Cre‑only controls; thus, floxed mice served as controls for all subsequent analyses. Experimental cohorts included mice 9–20 weeks of age, as specified in figure legends.

Glucose and insulin tolerance tests

For glucose tolerance tests (GTT), mice were fasted for 16 h before receiving 2 g/kg glucose orally or intraperitoneally. Blood glucose was measured from the tail vein at 0–120 min using a handheld glucometer (Contour Next One, Ascensia Diabetes Care). Insulin tolerance tests (ITT) were performed after a 2‑h fast, followed by intraperitoneal injection of 0.75 IU/kg insulin (Actrapid, Novo Nordisk) with sampling at 0–90 min.

Gastric emptying was assessed by acetaminophen absorption during an oral GTT using 0.1 g/kg acetaminophen (Merck), with plasma levels quantified using a commercial colorimetric assay kit (L3K assay kit, Sekisui Diagnostics).

Hormone measurements

Blood collected during tolerance tests was mixed with EDTA, diprotin A, and protease inhibitors and centrifuged for plasma isolation. Insulin, total GLP‑1, and total PYY were measured simultaneously using a U‑Plex multiplex assay (Meso Scale Diagnostics). For single‑analyte measurements, ELISA kits for insulin (Crystal Chem) and GLP‑1 (Meso Scale Diagnostics) were used. Ileal GLP‑1 content and islet insulin content were assessed from tissue homogenates using ELISA and normalized to protein content or tissue weight.

Physiological and metabolic phenotyping

Body weight and random‑fed glucose were measured weekly. Body composition was analysed by EchoMRI (Echo Medical Systems) and surface temperature by infrared imaging (FLIR Systems Inc.). Food and water intake were assessed following an overnight fast and measured at 4–48 h. Indirect calorimetry (OxyletPro Physiocage, Panlab) was used to record respiratory exchange, energy expenditure, locomotion, and feeding behaviour. For fast–refeed experiments, food was removed at ZT0 and reintroduced at ZT12. Measurements were performed at the Turku Centre for Disease Modelling (Finland).

Lymphocyte isolation and flow cytometry

Single‑cell suspensions from mesenteric lymph nodes (mLN), Peyer’s patches (PP), and intraepithelial lymphocytes (IELs) were prepared using established enzymatic and mechanical dissociation procedures (Qiu & Sheridan, 2018), followed by Percoll gradient separation for IELs. Cells were stained with viability dye and fluorochrome‑conjugated antibodies (Extended Data Table 2) and analysed using a CytoFLEX S cytometer (Beckman Coulter). Data were processed with FlowJo v10.8.1 (BD Life Sciences).

Cell culture

GLUTag (gifted by Daniel Drucker), MIN6 (gifted by Diana Toivola), and 293FT cells (gifted by Pekka Katajisto) were cultured under standard conditions in DMEM‑based media with heat‑inactivated FBS and antibiotics, using cell‑type‑specific supplements. EndoC‑βH1 cells were plated on ECM/fibronectin‑coated surfaces and maintained in Optiβ1 medium (Human Cell Design). All lines tested negative for mycoplasma contamination.

Murine intestinal organoids

Small intestinal crypts were generated using standard protocols (T. Sato et al., 2009). The composition of basal cell media was Advanced DMEM/F12 supplemented with antibiotics (50 U/ml penicillin, 50 µg/ml streptomycin), 10 mM HEPES, 2 mM GlutaMAX, 1 mM N-acetylcysteine, 1X N2 supplement, and 1X B27 supplement. The basal cell medium was supplemented with murine EGF (50 ng/ml), murine Noggin (100 ng/ml) and human R-spondin 1 (300 ng/ml) —referred to as ENR. Organoids were cultured in ENR medium and passaged every 5 days. Differentiation was induced by adding IWP‑2 (2 µM), DAPT (10 µM), or MEK inhibitor PD0325901 (1 µm). Stem cell enrichment used Wnt3a (100 ng/ml) and CHIR99021 (3 µM).

Mouse islet isolation

Pancreatic islets were isolated by collagenase P (Roche) digestion via the common bile duct, followed by density‑gradient separation and hand‑picking (Carter et al., 2009). Islets recovered overnight in RPMI‑1640 supplemented with 10% FBS.

Insulin and GLP-1 secretion assays

MIN6 and EndoC‑βH1 cells with modified Scinderin expression were incubated in basal buffer (Krebs-Ringer bicarbonate HEPES buffer or βKrebs buffer) before stimulation with glucose, KCl or IBMX. Secreted and intracellular insulin or proinsulin were quantified by ELISA (Mercodia) and normalized to protein content.

GLUTag cells and Scin‑deficient intestinal organoids were stimulated with glucose, L‑glutamine or forskolin/IBMX. Secreted GLP‑1 was quantified using a total GLP‑1 assay (Meso Scale Diagnostics) and normalized to protein or DNA content.

Chemical treatments

Actinomycin D (5 μg/mL, Merck) was used to inhibit transcription for mRNA stability assays. Brefeldin A (BFA, Merck) was applied to induce Golgi stress (16 h) or to acutely block anterograde trafficking (1 h) before immunofluorescence. Cell viability was assessed by MTT assay (Roche).

Modification of gene expression

Scin was knocked-down in MIN6 and EndoC-βH1 cells by transfection with small-interfering RNAs (siRNAs; 30 nM). Lipid-siRNA complexes were formed by mixing RNAiMAX lipofectamine (Thermo Fisher Scientific) with either non-targeting (NT) siRNA (#D-001810-10-05, Horizon) or Scin-targeting siRNA (#L-045307-01-0005, #L-015062-00-0005). RNA was collected 72 h post‑transfection; protein and functional assays were performed on day 4.

CRISPR–Cas9 knockout GLUTag cells were generated using lentiCRISPR v2 (Addgene) containing sgRNAs targeting *Scin* (sequence 5’-3’, KO1: CAGATGGACGACTATTTGGG, KO2: AAAAGACTTCTGCACGTGAA). Transduced cells were selected with puromycin and validated by immunoblotting.

GLUTag cells were transfected with plasmids encoding GFP‑tagged SAC1, SAC1‑K2A or SAC1‑LZ (a kind gift by Neale Ridgway); empty GFP vector served as control. Cells were analysed 48 h post‑transfection.

RNA extraction and qPCR

RNA was extracted using TRIzol (Thermo Fisher Scientific) or RNeasy Plus Micro (Qiagen) kits. cDNA synthesis used iScript (Bio-rad), and qPCR was performed with Ssofast Evagreen (Bio-Rad) on a CFX Opus 96 system (Bio-Rad). Expression levels were normalized to housekeeping genes using the 2‑ΔCt method. Primer sequences are listed in Extended Data Table 3.

Bioinformatic analysis

Publicly available single-cell RNA-seq data comprising healthy, prediabetic, and T2D pancreatic islet cells, were analyzed. UMI count matrices and metadata were processed in Seurat. Doublets were identified using scDblFinder, and cells with low feature counts (<200), high feature counts (>6,000), low UMI (<500), high UMI (>50,000), or high mitochondrial content (>20%) were excluded. Filtered datasets were normalized and variance-stabilized using SCTransform, regressing out mitochondrial gene content. The datasets were merged, retaining only the SCT assay, and 3,000 variable features were selected for integration. Principal component analysis (PCA) was performed on the SCT assay, and batch effects were corrected using Harmony. Cell neighbors and clusters were identified with FindNeighbors (dims 1–30) and FindClusters (resolution 0.4), and data were visualized using UMAP. To further resolve Beta cells, clusters with high expression of Beta cell markers (MAFA, NKX6-1, PDX1, IAPP, INS) were subset and reclustered following the same workflow. Aggregated signature gene scores were calculated using the AddModuleScore function in Seurat. Finally, pseudo-bulk samples were generated by summing counts of individual cells from the same cell type within each biological replicate, enabling assessment of transcriptomic changes at the bulk level.

For RNA sequencing, poly(A)‑selected libraries were sequenced on an Illumina NovaSeq X Plus (150‑bp paired‑end). Reads were quality filtered, mapped to GRCh38 or GRCm38 using HISAT2 and quantified as FPKM. Differential expression was performed using DESeq2 with Benjamini–Hochberg correction (adjusted P ≤ 0.05, |log2FC| ≥ 0.5). Functional enrichment was carried out using ClusterProfiler. Plots were generated using RStudio 4.5.2 (Posit). Additional datasets used for comparative analyses are listed in Extended Data Table 4.

Immunoblotting, subcellular fractionation and actin isolation

Cells and organoids were lysed directly in loading buffer, separated by SDS–PAGE or Tricine–SDS–PAGE (for GLP‑1 and proinsulin), and transferred to nitrocellulose or PVDF membranes. Immunoreactive bands were detected by infrared or chemiluminescence imaging and quantified using Fiji. Antibody information is provided in Extended Data Table 5.

Cytoplasmic fractions were isolated by sucrose‑gradient ultracentrifugation (0.45–2 M). Fractions were analysed by immunoblotting for organelle markers. F‑ and G‑actin fractions were separated by high‑speed centrifugation at 37°C and solubilized for immunoblotting.

Recombinant SCIN-HisTag was tested for PIP binding using differential scanning fluorimetry (Bio-Rad) and lipid strips (Echelon BioSciences) probed with anti‑HisTag antibody (Thermo Fisher Scientific).

Histology and microscopy

Tissues were fixed in formalin, paraffin‑embedded and sectioned (3–5 μm). H&E staining followed standard protocols. Tissue sections were deparaffinized, and heat-induced epitope retrieval was performed using Tris-EDTA buffer (pH 9) for immunostaining. The slides were blocked with blocking diluent (Vector laboratories) and incubated with primary and secondary antibodies. Antibodies used are listed in Extended Data Table 6. After antibody staining, the slides were counterstained with DAPI and mounted using ProLong Glass Antifade Mountant (Thermo Fisher Scientific). Negative controls lacked primary antibodies.

Cells were cultured on glass coverslips or cell culture chamber slides (Ibidi) for immunofluorescent staining. Upon reaching the appropriate confluence, cells were rinsed with PBS and fixed with 4% methanol-free formaldehyde for 10 minutes. Permeabilization was performed for 10 minutes using 0.2% Triton X-100. For blocking, PBS containing 0.05% Tween-20 and 1% BSA was used. Cells were then incubated sequentially with primary and secondary antibodies, followed by DAPI staining. The samples were mounted using ProLong or Ibidi mounting medium.

Transmission electron microscopy of isolated islets and ileum involved glutaraldehyde fixation, osmium postfixation, dehydration, embedding in Epon and ultrathin sectioning.

Images were acquired using confocal microscopy (Nikon A1R+), whole‑slide scanners (NanoZoomer S60 or Slide Strider scanner) and a transmission electron microscope (JEM‑1400). Image analyses used QuPath, Fiji, Huygens and associated plugins for segmentation, quantification, colocalization and morphometry.

Statistics

Data are shown as mean ± SEM (unless otherwise stated), and a p-value < 0.05 was considered statistically significant. Normality was assessed using the Shapiro–Wilk test and homoscedasticity using F‑test or Brown–Forsythe test. Statistical comparisons used unpaired two‑tailed Student’s t‑tests or Mann–Whitney U‑tests; one‑way ANOVA with Dunnett’s or Šidák’s correction or Kruskal–Wallis with Dunn’s test; and repeated‑measures two‑way ANOVA with Sidak correction for time‑course data. AUC was calculated by the trapezoidal method, and incremental AUC was baseline‑corrected. Categorical data were analysed by χ² test.
