## Supplemental Tables for "Scinderin-driven Golgi Actin Remodeling coordinates GLP-1 and insulin secretion to regulate glucose homeostasis"

**Supplemental Data Table 1**

Genotyping primers

| Name | Sequence FW (5’-3’) | Sequence RV (5’-3’) |
| --- | --- | --- |
| *Scin lox/lox* | GTTAGTATTCCTCACTGGCACCC | ATGTTTCAGGACAGGAGTCTGAGC |
| *Scin Δ/Δ* | GACGACTATTTGGGTGGCAAG,  TCACGCGTATAACTTCGTATAGCA | GCAGGGTTTCTGAGGCACAT |
| *Vil1‑Cre* | GCCTTCTCCTCTAGGCTCGT | TATAGGGCAGAGCTGGAGGA,  AGGCAAATTTTGGTGTACGG |
| *Ins1‑Cre* | GTCAAACAGCATCTTTGTGGTC, GCTGGAAGATGGCGATTAGC | GGAAGCAGAATTCCAGATACTTG |

**Supplemental Data Table 2**

List of antibodies used in flow cytometry experiments

| Marker | Fluorochrome | Clone | Manufacturer | Catalog # | Target cell |
| --- | --- | --- | --- | --- | --- |
| TCR γδ | eFluor450 | GL3 | eBioscience | 48-5711-82 | γδ T lymphocyte |
| Viability dye | FVS510 | - | BD Biosciences | 564406 | Viable cells |
| CD45 | FITC | 30-F11 | BD Biosciences | 553079 | Leukocyte |
| CD8α | PerCP-Cy5.5 | 53-6.7 | BD Biosciences | 551162 | Cytotoxic T cell |
| TCR β | PE | H57-597 | eBioscience | 12-5961-81 | αβ T lymphocyte |
| CD3 | PE-Cy7 | 17A2 | BD Biosciences | 560591 | T lymphocyte |
| CD4 | APC-eFluor780 | GK1.5 | eBioscience | 47-0041-80 | Helper T cell |
| CD19 | Super Bright 780 | 1D3 | eBioscience | 78-0193-82 | B lymphocyte |

**Supplemental Data Table 3**

Sequence of primers used in qPCR

| Gene | Species | Forward primer 5' - 3' | Reverse primer 5' - 3' |
| --- | --- | --- | --- |
| *Scin* | Mouse | TGGCGAAGCTCTACATGGTTT | TCCTCTCCTGTGGGTTAGCA |
| *Gcg* | Mouse | GTGACTGGCACGAGATGTTG | CTTCCCAGAAGAAGTCGCCA |
| *Chga* | Mouse | GTGCGTCCTGGAAGTCATCTCC | GAGAGCCAGGTCTTGAAGTTCC |
| *Pdx1* | Mouse | CCGAATGGAACCGAGCCTG | GCAGTACGGGTCCTCTTGTT |
| *Rpl32* | Mouse | CAATGTGTCCTCCTCTAAGAACCGAAA | CCTGGCGTTGGGATTGG |
| *Gapdh* | Mouse | TGTGTCCGTCGTGGATCTGA | CCTGCTTCACCACCTTCTTGA |
| *rRNA18s* | M/H | GTAACCCGTTGAACCCCATT | CCATCCAATCGGTAGTAGCG |
| *SCIN* | Human | CTGGAGGATTGAGAAGCTGGAG | GGAACACTCCTTTCCGAGCCA |
| *PPIA* | Human | ATGGCAAATGCTGGACCCAACA | ACATGCTTGCCATCCAACCACT |

**Supplemental Data Table 4**

Datasets used in transcriptomic analyses

| Dataset type | Source / Accession |
| --- | --- |
| Mouse intestinal epithelium scRNA‑seq | Haber et al., 2017; Single Cell Portal |
| Human ileum + EE organoids scRNA‑seq | GSE146799 |
| GLUTag RNA‑seq | GSE193866 |
| Mouse intestinal organoid ChIP‑seq & GRO‑seq | GSE78761 |
| Human islet ATAC‑seq & RNA‑seq | Islet Regulome Browser |
| Human islet snRNA‑seq | Mummey et al., 2024 |
| Human pancreas development scRNA‑seq | Olaniru et al., 2023 |
| Deep murine islet proteome | PXD024522 |
| EndoC‑βH1 transcriptome | GSE118588 |
| Human islet BFA‑treated RNA‑seq | GSE152615 |
| RNA-seq SC-islets differentiation | Balboa et al., 2022 |
| ATAC-seq SC-islets differentiation | Alvarez-Dominguez et al., 2020 |
| ATAC-seq beta, alpha cells, adult tissues | Zhang et al., 2021 |
| Type-1 diabetes pancreatic islets RNAseq | GSE228267 |
| Type-2 diabetes pancreatic islets scRNAseq | GSE221156 |
| H3K27me3 ChIPseq islets | GSE224061 |

**Supplemental Data Table 5**

List of antibodies used in immunoblot experiments

| Target | Dilution | Manufacturer | Catalog # |
| --- | --- | --- | --- |
| SCIN | 1:1000 | abcam | ab199723 |
| (PRO)INS | 1:1000 | Cell Signaling | 8138 |
| GLP-1 | 1:1000 | abcam | ab23472 |
| GM130 | 1:1000 | BD Biosciences | 610822 |
| TGN38 | 1:1000 | Santa Cruz Biotechonology | sc-166594 |
| β-tubulin | 1:20,000 | Merck | T7816 |
| H3 | 1:35,000 | abcam | ab1791 |
| Actin | 1:20,000 | Merck | MAB1501R |
| GAPDH | 1:1000 | abcam | ab8245 |
| CALR | 1:1000 | Novus Biologicals | NBP1-47518 |
| anti-mouse-HRP | 1:1000 | Agilent (Dako) | P0260 |
| anti-rabbit-HRP | 1:1700 | Agilent (Dako) | P0217 |
| anti-mouse-IRDye 800CW | 1:20,000 | LI-COR | 926-32210 |
| anti-rabbit-IRDye 680RD | 1:20,000 | LI-COR | 926-68071 |

**Supplemental Data Table 6**

List of antibodies used in immunofluorescent staining

| Target | Dilution | Manufacturer | Catalog # |
| --- | --- | --- | --- |
| SCIN | 1:250 | abcam | ab199723 |
| SCIN | 1:100 | Atlas antibodies | HPA020518 |
| SCIN | 1:100 | Santa Cruz Biotechonology | sc-376136 |
| PROINS | 1:50 | R&D systems | MAB13361 |
| INS | Undiluted | Agilent (Dako) | IR002 |
| GCG | 1:2000 | Merck | G2654 |
| GCG | 1:200 | Cell Signaling | 2760 |
| SST | 1:100 | Merck | MAB354 |
| PPY | 1:100 | Santa Cruz Biotechonology | sc-514155 |
| GLP-1 | 1:1000 | abcam | ab23472 |
| PYY | 1:400 | abcam | ab22663 |
| GIP | 1:100 | abcam | ab209792 |
| 5-HT | 1:500 | abcam | ab66047 |
| CCK | 1:100 | Thermo Fisher Scientific | PA5-103116 |
| MUC2 | 1:100 | Santa Cruz Biotechonology | sc-15334 |
| KI-67 | 1:250 | Thermo Fisher Scientific | MA5-14520 |
| CC3 | 1:400 | Cell Signaling | 9661 |
| GFP | 1:100 | Origene | AB0020-200 |
| GM130 | 1:200 | BD Biosciences | 610822 |
| CALR | 1:200 | Novus Biologicals | NBP1-47518 |
| Phalloidin-488 | 1:1000 | abcam | ab176753 |
| Phalloidin-568 | 1:400 | Invitrogen | A12380 |
| GSN | 1:50 | Santa Cruz Biotechonology | sc-514502 |
| GRASP65 | 1:200 | Thermo Fisher Scientific | PA3-910 |
| ERGIC-53 | 1:50 | Santa Cruz Biotechonology | sc-365158 |
